# Spatial control of sensory adaptation modulates mechanosensing in *Pseudomonas aeruginosa*

**DOI:** 10.1101/2024.02.27.582188

**Authors:** Ramiro Patino, Marco J. Kühn, Henriette Macmillan, Yuki F. Inclan, Ivan Chavez, John Von Dollen, Jeffrey R. Johnson, Danielle L. Swaney, Nevan J. Krogan, Alexandre Persat, Joanne N. Engel

## Abstract

Sensory signaling pathways use adaptation to dynamically respond to changes in their environment. Here, we report the mechanism of sensory adaptation in the Pil-Chp mechanosensory system, which the important human pathogen *Pseudomonas aeruginosa* uses to sense mechanical stimuli during surface exploration. Using biochemistry, genetics, and cell biology, we discovered that the enzymes responsible for adaptation, a methyltransferase and a methylesterase, are segregated to opposing cell poles as *P. aeruginosa* explore surfaces. By coordinating the localization of both enzymes, we found that the Pil-Chp response regulators influence local receptor methylation, the molecular basis of bacterial sensory adaptation. We propose a model in which adaptation during mechanosensing spatially resets local receptor methylation, and thus Pil-Chp signaling, to modulate the pathway outputs, which are involved in *P. aeruginosa* virulence. Despite decades of bacterial sensory adaptation studies, our work has uncovered an unrecognized mechanism that bacteria use to achieve adaptation to sensory stimuli.

## Introduction

Living systems must adapt to ever changing external signals to elicit proper biochemical responses^1^. For example, neurons rapidly respond to changes in the environment by adapting to a broad range of temperatures or smells^2^. The biological process of maintaining sensitivity in different background states is known as sensory adaptation and is a general feature of sensory signaling pathways in all domains of life^3^.

Much of what we know about sensory adaptation in bacteria comes from extensive studies of the *Escherichia coli* chemotaxis pathway, which has served as a model system. This sensory pathway responds to and signals across several orders of magnitude of ligand concentrations without reaching saturation^4,5^. By primarily binding chemical ligands^6,7^, methyl-accepting chemotaxis proteins (MCPs, also known as chemoreceptors) signal to downstream components that promote a chemotactic response^8^. MCPs activate a histidine kinase (CheA) that can transfer a phosphoryl group to a response regulator (CheY)^9,10^. Upon phosphorylation, CheY binds to the base of the flagellar motor to change the direction of flagellar rotation, thereby controlling swimming direction^11^.

Sensory adaptation is achieved through a negative feedback loop that involves reversible methylation of MCPs^12,13^. MCP methylation is catalyzed by the constitutively active CheR methyltransferase^14–16^, which enhances MCP signaling activity^17^. Activated by CheA-dependent phosphorylation, like CheY, the CheB methylesterase demethylates MCPs, thereby decreasing MCP signaling activity^18,19^. By tuning MCP sensitivity and signaling activity, MCP methylation therefore serves as the molecular basis of bacterial sensory adaptation. Overall, sensory adaptation ensures that *E. coli* can robustly chemotax in response to small changes in ligand concentration that occur in chemical gradients ^20–22^.

*Pseudomonas aeruginosa*, an important human pathogen, relies on four sensory signaling pathways to adapt to its environment, including during human infections^23^. Two sensory pathways are involved in *P. aeruginosa* chemotaxis, while a third sensory pathway, the Wsp system, participates in biofilm formation^23–25^. We have recently shown that the Pil-Chp system, the fourth sensory pathway, senses mechanical stimuli generated by surface contact, thereby participating in mechanosensing^26^. Although all four of the sensory pathways encode predicted sensory adaptation enzymes, minimal information is available about how sensory adaptation functions during *P. aeruginosa* mechanosensing.

We have demonstrated that the Pil-Chp system responds to mechanical signals transmitted by type IV pili (TFP) upon surface contact^26^. TFP are polarly localized protein filaments that undergo repeated cycles of extension, attachment, and retraction ^27^. By localizing TFP to one cell pole (i.e. leading pole), cells propel themselves forward in a specialized form of surface exploration called twitching motility^27^. Surface contact through TFP stimulates PilJ^26,28^, the Pil-Chp MCP, initiating signaling through the downstream Pil-Chp components to trigger two outputs: the polarized deployment of additional TFP and the production of the second messenger, cyclic adenosine monophosphate (cAMP)^26,29–32^.

Through deploying more TFP at one cell pole, *P. aeruginosa* can establish a local positive feedback loop that promotes twitching in the direction of mechanical stimulation (“mechanotax”) over long distances and activate a cAMP-dependent acute virulence program. Either spontaneously or upon mechanical perturbations, such as cell-cell collisions, the Pil-Chp system also inverts TFP polarity to induce reversals in twitching motility direction^33,34^. The twitching motility reversals are coordinated by the spatial organization of PilG and PilH, two antagonistic CheY-like response regulators that are downstream components of the Pil-Chp system^34^. Local Pil-Chp activation at the leading pole recruits and activates PilG through phosphorylation, which drives the positive feedback loop that sustains forward movement. Accumulation of bipolarly localized PilH serves as a brake to inhibit PilG phosphorylation and enable twitching motility reversals.

The Pil-Chp regulon also encodes two predicted sensory adaptation enzymes, the methyltransferase PilK and the methylesterase ChpB^31,32,35^. However, if and how sensory adaptation takes place in the Pil-Chp system during mechanosensing is unresolved. Here, we test the hypothesis that PilK and ChpB are bona fide sensory adaptation enzymes that control PilJ methylation and sensory adaptation during mechanosensing. We used PilK and ChpB mutants in conjunction with a method to assess PilJ methylation *in vivo* to demonstrate that PilJ undergoes reversible methylation that modulates Pil-Chp signaling. Using functional fluorescent protein fusions, we discovered that the Pil-Chp sensory adaptation enzymes are segregated to opposite cell poles in twitching *P. aeruginosa* cells. We further demonstrate that PilG and PilH dictate the localization behavior of PilK and ChpB, which is critical to control the PilJ methylation state. Our study thus reveals the mechanism of sensory adaptation during bacterial mechanosensing.

## Results

### PilK and ChpB modulate twitching motility reversals and cAMP production

In addition to encoding two antagonistic response regulators, PilG and PilH, the Pil-Chp operon encodes two predicted sensory adaptation enzymes, the PilK methyltransferase and ChpB methylesterase. While the response regulators have antagonistic effects on the physiological outcomes of Pil-Chp signaling (Figure 1A), limited information exists about how PilK and ChpB impact the system’s two outputs: twitching motility and cAMP production^31,32^.

**Figure 1.**
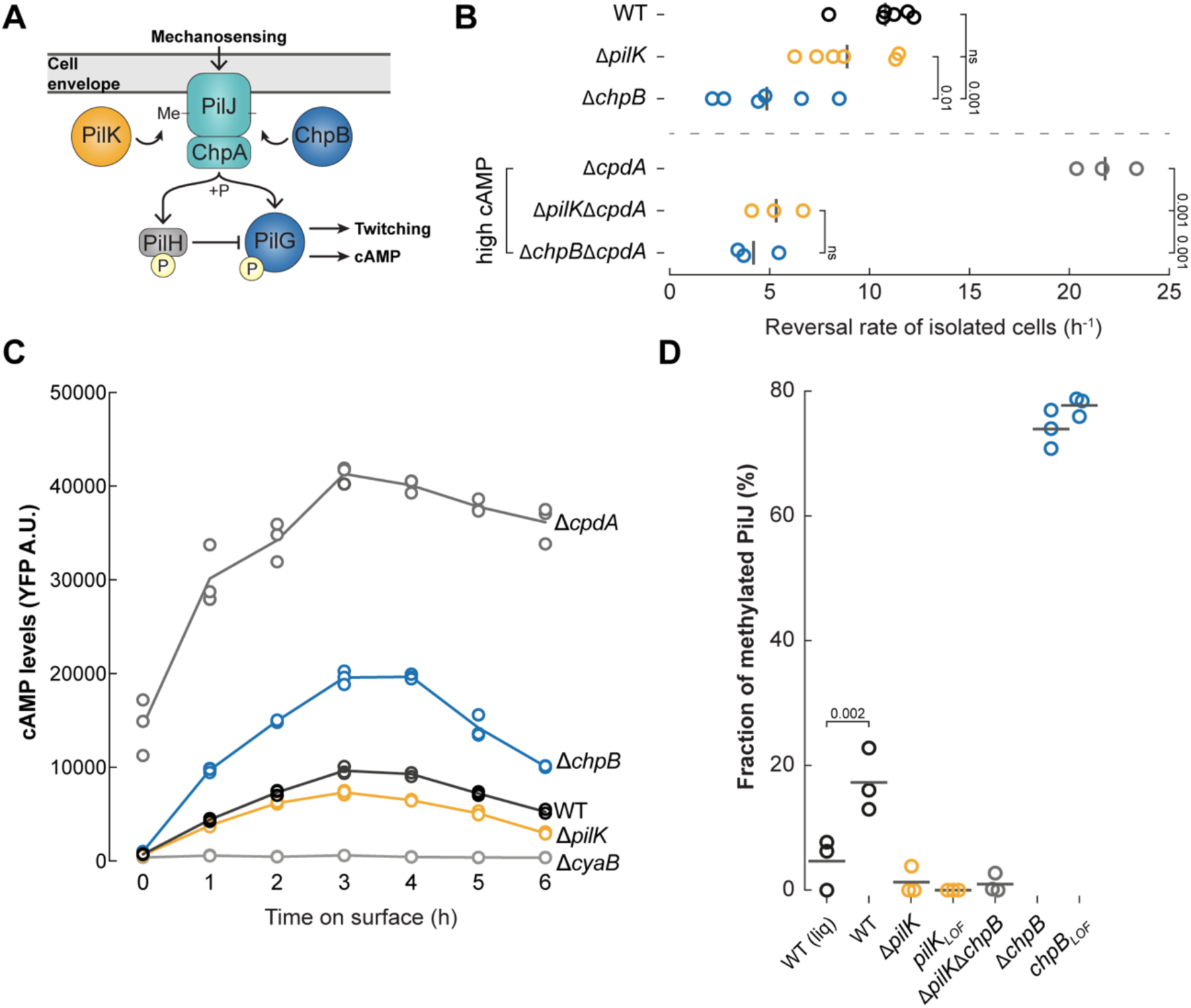
ChpB and PilK modulate twitching reversals and cAMP production by controlling PilJ methylation. **(A)** Schematic of the Pil-Chp mechanosensory system. Mechanical stimuli are sensed by the MCP PilJ, which stimulates ChpA autophosphorylation. The phosphoryl group is transferred to PilG. PilG∼P stimulates twitching motility and cAMP production. PilH∼P inhibits PilG. The methyltransferase PilK and the methylesterase ChpB control PilJ methylation. **(B)** PilK and ChpB modulate twitching motility reversal rates. Shown are the spontaneous reversal rates of isolated motile cells after 2 h surface growth in WT background (upper panel) or *ΔcpdA* (lower panel, with elevated cAMP levels). Circles, median of each biological replicate. Vertical bars, mean across biological replicates. Vertical numbers indicate p-values (ANOVA and Tukey’s post hoc test; ns, not significant). For corresponding reversal rates after cell-cell collisions see Suppl Fig 2A. **(C)** PilK and ChpB fine-tune the amplitude of surface-dependent cAMP production over time. cAMP was measured with a transcriptional reporter by flow cytometry at each indicated time point for the indicated strains. A representative graph of two independent experiments, with three biological replicates per experiment, is shown. Circles, median YFP fluorescence of each biological replicate (>30,000 cells each). Each line connects the means of biological replicates. A.U., arbitrary units. **(D)** PilJ methylation *in vivo*. Shown is the fraction of methylated PilJ (methylated PilJ signal/total PilJ signal) of the indicated strains expressing chromosomal 3xFLAG-PilJ. Circles, biological replicates. Horizontal bar, mean across biological replicates. P-value shown for the WT liquid vs solid comparison (ANOVA with Tukey’s post hoc test). LOF, loss of function. For corresponding twitching motility assays, cAMP assays, and immunoblot images see Suppl Figs 1A, 1C, and Suppl Fig 3A, respectively.

Although deletion of *pilK* and *chpB* have minimal effects on twitching motility as assayed by the subsurface stab assay ^31,32,35^(Suppl Fig 1A), whether PilK and ChpB alter single cell twitching motility has not been explored. To address this question, we calculated the reversal frequencies of isolated Δ*pilK* and Δ*chpB* twitching cells from movies, either upon spontaneous reversals or upon collision with other bacteria^33,34^. Isolated motile cells of either Δ*pilK* or Δ*chpB* exhibited slightly lower reversal frequencies than WT, with only Δ*chpB* reaching statistical significance (Fig 1B; Suppl Fig 2A). To further explore potential differences in the reversal rates, we examined whether the decrease in reversal frequencies observed in Δ*pilK* and Δ*chpB* were amplified in a sensitized background with elevated cAMP (*ΔcpdA*). In the absence of CpdA, *P. aeruginosa* has constitutively elevated cAMP, with a concomitant increase in the number of TFP and reversal frequency^31,33,36,37^. The reversal frequency of either *ΔpilKΔcpdA* or *ΔchpBΔcpdA* was significantly decreased compared to *ΔcpdA* (Fig 1B; Suppl Fig 2A). The decrease in reversal frequencies was not due to altered twitching motility speed, as the twitching motility speed of *ΔpilKΔcpdA* and *ΔchpBΔcpdA* was not statistically different compared to *ΔcpdA* (Suppl Fig 2B). We conclude that PilK and ChpB modulate the frequency of twitching motility reversals, especially when more mechanically active TFP are present.

Activation of the Pil-Chp system by mechanosensing also promotes cAMP production, albeit on the timescale of hours^26,29^. cAMP binds to the Vfr transcriptional activator to upregulate the transcription of >200 genes, including an acute virulence program^38,39^. We thus used Δ*pilK* and Δ*chpB* strains to examine the role of these sensory adaptation enzymes in cAMP production over time (Fig 1C). We quantified cAMP levels from liquid-grown bacteria and from bacteria plated for up to 6 h on an agar surface, where mechanosensing occurs. In WT, cAMP levels increased upon surface contact. Maximal cAMP levels were observed at 2-3 h of surface growth and then decreased towards baseline levels at ∼ 6 h of surface growth, consistent with our previously reported results^26^. As expected, cAMP levels were constitutively low in *ΔcyaB* (deletion of the primary adenylate cyclase)^26,31,38^ and persistently elevated in *ΔcpdA* (deletion of the cAMP-specific phosphodiesterase)^26,36,37^. In *ΔpilK*, cAMP levels increased upon surface contact and decreased towards baseline, similar to WT, but the peak cAMP level was slightly decreased (Fig 1C). In contrast, cAMP levels were elevated at all timepoints in *ΔchpB* as compared to WT. However, cAMP levels still decreased towards baseline levels over time without ChpB (Fig 1C). Although sensory adaptation enzymes typically reset the physiological outcome(s) of bacterial sensory systems, our data suggest that PilK and ChpB are not required to return cAMP levels back towards pre-stimulation levels. Rather, resetting cAMP levels to the unstimulated state (i.e., liquid growth) requires the phosphodiesterase CpdA (Fig 1C), whose transcription is regulated by cAMP^37^. We suggest instead that PilK and ChpB fine-tune cAMP production as well as reversal rates during twitching motility, likely by controlling the methylation state of the Pil-Chp receptor.

### PilJ methylation increases in response to mechanosensing and is controlled by PilK and ChpB

PilJ, the MCP of the Pil-Chp system, is required to sense and to respond to mechanical inputs generated during surface contact^26,29^; however, whether PilJ becomes methylated *in vivo* by the action of PilK and ChpB has never been demonstrated. We therefore tested whether PilJ methylation is detectable *in vivo* and whether PilJ methylation changes during liquid growth compared to surface growth. For these studies, the native *pilJ* gene was replaced with a *3xFlag-pilJ* fusion for detection by immunoblot analysis. Full-length 3xFlag-PilJ migrated as a single band of the expected molecular weight on conventional SDS-PAGE (Suppl Fig 1B). The chromosomally integrated 3xFlag-PilJ is functional, as demonstrated by near wild-type (WT) levels of subsurface twitching motility (Suppl Fig 1A) and cAMP production (Suppl Fig 1C). We used a well-established low-bis SDS-PAGE separation protocol to detect and quantify MCP methylation *in vivo* (hereafter referred to as methylation immunoblots)^40–45^. We detected two differentially migrating species of PilJ during liquid and surface growth (Suppl Fig 3A). By analogy to the migration of *E. coli* MCPs in methylation immunoblots^40–45^, the slower migrating band is predicted to represent the unmethylated PilJ state, and the faster migrating band is predicted to represent a methylated form of PilJ (which we validate below). By calculating the fraction of methylated PilJ, we demonstrated that methylated PilJ was significantly increased after 2 h of surface growth compared to liquid growth (p=0.002; Fig 1D), suggesting that PilJ methylation increases in response to mechanical stimuli generated during surface contact.

Since surface contact promotes cAMP synthesis^26,29^ (Fig 1C), and many TFP and Pil-Chp genes are positively regulated by cAMP^38^, we assessed whether the increase in PilJ methylation after surface growth was a secondary consequence of increased cAMP levels. The fraction of methylated PilJ did not significantly change in strains with low cAMP levels (*ΔcyaB*) or with high cAMP levels (*ΔcpdA*; Suppl Fig 3B,C). We conclude that the increase in PilJ methylation after surface contact is likely a direct response to mechanosensing, rather than a secondary effect from the increase in cAMP levels.

We next tested whether PilK and ChpB control the methylation state of PilJ and to definitively assign methylation states to the two PilJ species observed in the methylation immunoblots. We introduced *3xFlag-pilJ* into Δ*pilK*, *ΔchpB*, and *ΔpilKΔchpB* (Fig 1D; Suppl Fig 3A). In methylation immunoblots of Δ*pilK and ΔpilKΔchpB*, we primarily observed a slower migrating band, which likely corresponds to unmethylated PilJ, and an almost undetectable faction of methylated PilJ. In *ΔchpB*, the faster-migrating 3xFlag-PilJ band was the most abundant species, likely corresponding to methylated PilJ, with a corresponding increase in the fraction of methylated PilJ. Thus, like the migration behavior of *E. coli* MCPs^40–45^, methylated PilJ migrates faster than unmethylated PilJ in this experimental setup.

To link PilK and ChpB function to their enzymatic activity, we replaced the corresponding WT *pilK* and *chpB* alleles in *3xFlag-pilJ* with the respective catalytic site mutants, based on homology to *E. coli* CheR and CheB, respectively^15,46^ (*pilK*_R92E_ and *chpB*_H184Y,_ hereafter referred to as *pilK_LO_*_F_ and *chpB_LOF_*). Twitching motility subsurface stab assays (Suppl Fig 1A), cAMP measurements (Suppl Fig 1C), and methylation immunoblots (Fig 1D) showed that each loss-of-function (LOF) mutant phenocopied the respective *pilK* and *chpB* deletion mutant. We conclude that the enzymatic activity of PilK and of ChpB is needed to control PilJ methylation, thereby validating that PilK and ChpB are bona fide bacterial adaptation enzymes. Overall, we propose that PilK and ChpB modulate twitching motility and cAMP production by controlling PilJ methylation in response to mechanosensing.

### PilK and ChpB are spatially segregated to opposite poles in twitching cells

We have shown that the dynamic localization of PilG and PilH, the two response regulators of the Pil-Chp system, establish TFP polarity and dictate reversals^34^. We thus considered whether PilK and ChpB might also exhibit dynamic subcellular localization that could explain the mechanism by which they control PilJ methylation and modulate outcomes of Pil-Chp signaling. To test this hypothesis, we quantified static population-averaged fluorescent profiles of chromosomal ChpB-mNeonGreen (ChpB-mNG) and plasmid-expressed mNG-PilK (Fig 2). From the fluorescence profiles, we derived (i) a polar localization index, measuring the ratio of polar versus cytoplasmic localization and (ii) an asymmetry index, which quantifies the fluorescence intensity difference between the two poles^33,34^.

**Figure 2.**
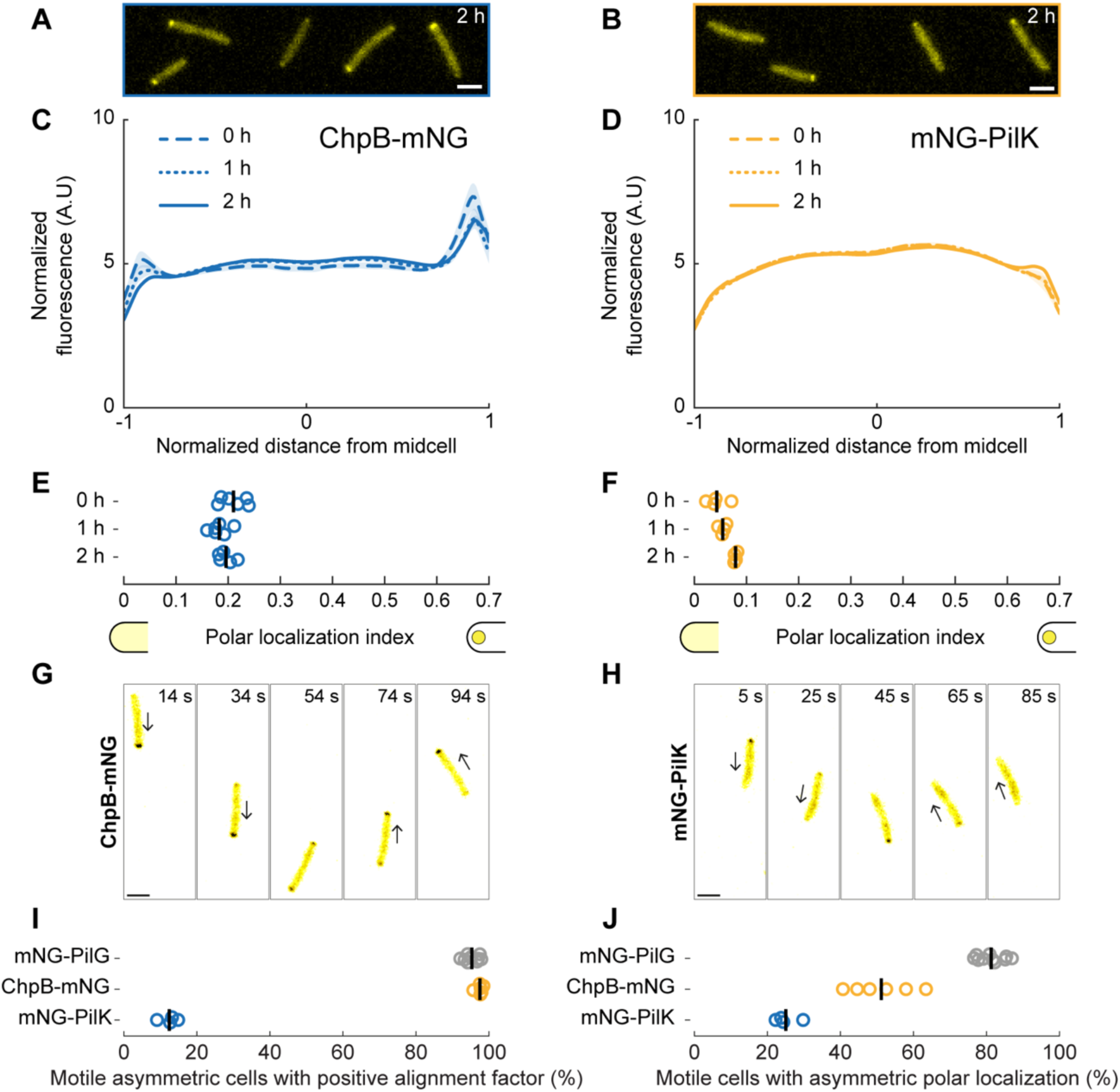
PilK and ChpB localize to the lagging and leading cell poles, respectively, of twitching *P. aeruginosa* cells. **(A, B)** Representative fluorescence snapshots of motile cells expressing **(A)** chromosomal ChpB-mNG or **(B)** plasmid-expressed mNG-PilK in Δ*pilK* after 2 h of surface growth. Scale bar, 2 µm. **(C, D)** Time course of normalized average fluorescence profiles for **(C)** ChpB-mNG or **(D)** mNG-PilK. The length of each cell is normalized so that the dim pole is positioned at *x* = −1, the bright pole at *x* = 1. The fluorescent profile for each cell is normalized by the total fluorescence. Solid lines, mean normalized fluorescence profiles across replicates. Shaded area, standard deviation across replicates. A.U., arbitrary units. **(E, F)** Corresponding polar localization index measuring the relative fraction of the fluorescence signal at the poles compared to the cytoplasm. Circles, median of each biological replicate. Vertical bars, mean across biological replicates **(G, H)** Time-lapse fluorescent snapshots of reversing cells taken from Movies 1 and 2. **(G)** ChpB-mNG localizes to the leading cell pole. **(H)** mNG-PilK localizes to the lagging cell pole. Both proteins switch poles during the twitching motility reversal. Arrows indicate direction of twitching motility. **(I)** Percent of motile cells with asymmetric polar localization that are twitching in the direction of their brightest pole. In most motile cells expressing ChpB-mNG, the brightest fluorescence localizes to the leading cell pole (alignment >0), similar to mNG-PilG^33,34^. Most motile cells expressing mNG-PilK have the brightest fluorescence at the lagging cell pole (alignment <0). Circles, median of each biological replicate. Vertical bars, mean across biological replicates. **(J)** Percent of motile cells with asymmetric polar localization of the indicated fluorescent fusion (intensity of the dim pole ≤ 80 % compared to the bright pole). Circles, median of each biological replicate. Vertical bars, mean across biological replicates. For corresponding asymmetry index and mean cell fluorescence measurements see Suppl Fig 5A.

We verified that ChpB-mNG was functional, as evidenced by WT twitching motility in sub-surface stab assays, near WT cAMP production, and expression of a full-length protein as assessed by immunoblot (Suppl Fig 4). We observed polar foci of ChpB-mNG from the moment cells encounter the surface (time = 0 h) to ∼2 h of surface contact (Fig 2A, C, E; Suppl Fig 5A; later time points were not examined). In most cells, the fluorescent signal appeared asymmetrically localized, with a more pronounced peak in the fluorescent profile defining the bright pole (close to *x* = 1; Fig 2C; Suppl Fig 5A). Since chromosomally integrated PilK fusion strains exhibited decreased twitching motility (Suppl Fig 6A), we instead expressed mNG-PilK at low levels from a plasmid in a Δ*pilK* background. Plasmid-expressed mNG-PilK exhibited WT twitching motility and restored PilJ methylation in Δ*pilK* (Suppl Fig 6B, C), although no full-length protein species could be detected by immunoblotting (Suppl Fig 6D, E). In contrast to chromosomal ChpB-mNG, mNG-PilK polar foci were very dim upon initial surface contact (time = 0 h) but became more visible after 2 h of surface growth, with the fluorescent signal appearing primarily at one cell pole (Fig 2B, D, F; Suppl Fig 5A). We observed that low cAMP levels (*ΔcyaB)* diminished polar localization of mNG-PilK and ChpB-mNG, while elevated cAMP levels (*ΔcpdA)* enhanced their polar localization (Suppl Fig 7). These results suggest that cAMP levels influence the polar localization of PilK and ChpB, without influencing the fraction of methylated PilJ (Suppl Fig 3C).

As both PilK and ChpB exhibit asymmetric polar localization, we determined whether they preferentially localize to the leading or to the lagging pole of twitching cells. For this assessment, we calculated an alignment factor between twitching direction and fluorescence asymmetry. A positive alignment factor indicates localization to the leading pole, whereas a negative alignment factor indicates localization to the lagging pole. Similar to mNG-PilG^33^, ChpB-mNG exhibited a positive alignment factor in almost all motile cells, demonstrating preferential localization to the leading pole in twitching cells (Figs 2G, I, J). In contrast, mNG-PilK exhibited a negative alignment factor in most motile cells, showing preferential localization to the lagging pole in twitching cells (Figs 2H, I, J). We also observed that the localization of mNG-PilK and ChpB-mNG was dynamic. Upon a twitching motility reversal, mNG-PilK and ChpB-mNG fluorescent foci changed polarity to the new lagging and new leading pole, respectively (Figs 2G, H; Movie 1 for PilK and Movie 2 for ChpB). In summary, mNG-PilK and ChpB-mNG are spatially segregated to opposite poles in twitching *P. aeruginosa* cells. By primarily localizing to the leading pole, where mechanosensing takes place, ChpB may demethylate and decrease PilJ signaling. At the opposite (lagging) pole, PilK may methylate and prime PilJ for efficient signaling once mechanosensing begins at the new leading pole.

### The response regulators PilG and PilH control the PilK and ChpB localization patterns

To understand how PilK and ChpB are spatially segregated to opposite cell poles in twitching cells, we investigated the mechanisms specifying their localization patterns. As both PilG^33,34^ and ChpB (Fig 2) dynamically localize to the leading pole, we first tested whether ChpB is required for PilG asymmetric polar localization. Polar localization and asymmetry indices of mNG-PilG were unchanged in *ΔchpB* (Suppl Fig 9), indicating that PilG localization is not regulated by ChpB. Remarkably, polar localization of ChpB-mNG was lost in *ΔpilG,* resulting in a polar localization index of 0, both in high (Figs 3A, C, E) and low cAMP backgrounds (Suppl Fig 10). Moreover, ChpB asymmetric polar localization was enhanced in *ΔpilH,* in a similar manner to PilG^34^ (Fig 3A, C, E, G), while ChpB did not play a role in the localization of chromosomally expressed and functional mNG-PilH (Suppl Fig 9). Polar localization of ChpB-mNG was lost in *ΔpilGΔpilH* (Fig 3A, C, E), demonstrating that PilH indirectly regulates ChpB asymmetric polar localization through PilG. Overall, these data suggest that PilG directs ChpB to the leading pole. Given that ChpB asymmetric polar localization depends on PilG, we considered whether PilG could recruit ChpB by direct or indirect binding. We performed affinity-purification mass spectrometry on lysates prepared from bacteria transformed with a plasmid overexpressing PilG-HA as the bait, PilH-HA as a comparison bait, or from untransformed WT as a control. Protein-protein interactions were scored by SAINT from a scale of 0 to 1^47^. ChpB was significantly enriched in affinity purified PilG-HA lysates (SAINT score 0.98), while no ChpB peptides were detectable in affinity purified PilH-HA (Supplemental Table 1). Therefore, a protein-protein interaction between PilG and ChpB may account for their linked localization patterns.

**Figure 3.**
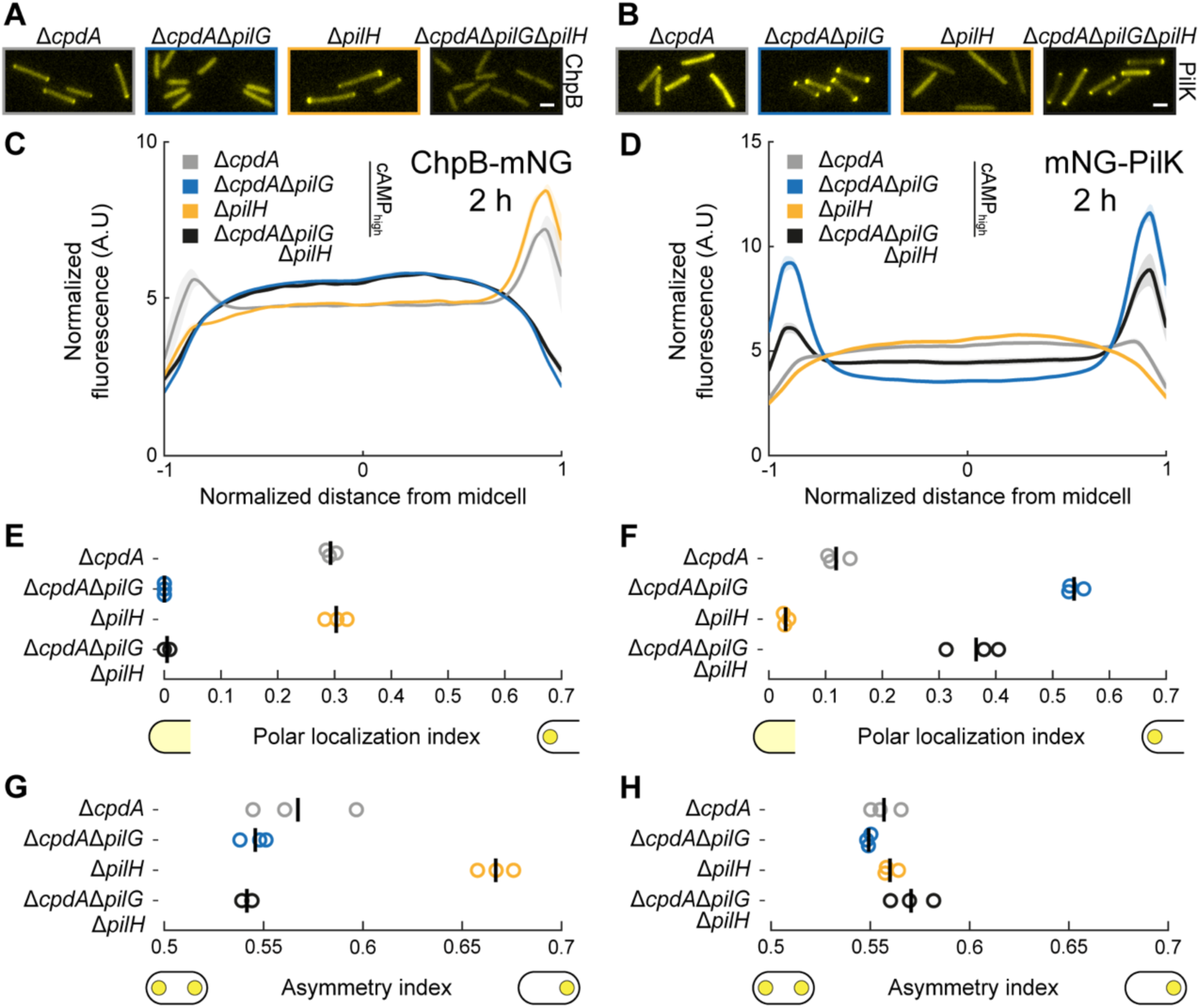
The response regulators PilG and PilH control the localization patterns of PilK and ChpB. **(A, B)** Representative fluorescence snapshots of motile cells expressing **(A)** chromosomal ChpB-mNG or **(B)** plasmid-expressed mNG-PilK in Δ*pilK* after 2 h of surface growth. Scale bar, 2 µm. **(C, D)** Fluorescence profiles of **(C)** ChpB-mNG or **(D)** mNG-PilK in Δ*pilG* and in Δ*pilH* mutants after 2 h of surface growth. *cpdA* was deleted in *pilG* mutants to eliminate potential confounding effects of low cAMP levels. *ΔpilH* has elevated cAMP. Solid lines, mean normalized fluorescence profiles across biological replicates. Shaded area, standard deviation across biological replicates. A.U., arbitrary units. **(E-H)** Corresponding measurements of polar localization index and asymmetry index. Circles, median of each biological replicate. Vertical bars, mean across biological replicates. For corresponding mean cell fluorescence measurements see Suppl Fig 5B.

We also investigated factors regulating PilK localization to the lagging pole. Since ChpB primarily localizes to the leading pole, we tested whether ChpB excludes PilK from the leading pole. The polar localization index of mNG-PilK was slightly increased without *chpB*; however, this increase in polar localization was likely due to the high cAMP levels in *ΔchpB*, as a similar increase was observed in *ΔcpdA* (Suppl Fig 11A, C, E). These data suggest that mNG-PilK can localize to the lagging pole without *chpB*. In a similar trend, ChpB-mNG localization was unchanged in *ΔpilK* (Suppl Fig 11B, D, F, H). Thus, PilK and ChpB localization patterns are largely independent. We next tested whether the PilK polar localization is regulated by the Pil-Chp response regulators, as is the case with ChpB. The polar localization of mNG-PilK was dramatically increased in *ΔpilG* and almost undetectable in *ΔpilH*, independent of cAMP levels (Figs 3B, D, F; Suppl Fig. 10B, D, F). However, as mNG-PilK shows prominent polar localization in *ΔpilGΔpilH* (Fig 3B, D, F), our data suggest that PilK can localize to the lagging pole independently of PilG and PilH. We propose that the Pil-Chp response regulators instead ensure that PilK is segregated to the lagging pole. PilG may prevent PilK accumulation at the leading pole while PilH allows PilK localization at lagging pole in a PilG-dependent manner. Taken together, these findings indicate that PilG and PilH establish the subcellular architecture of the Pil-Chp sensory adaptation enzymes.

### The ChpA histidine kinase and PilH activation affect ChpB localization

We previously demonstrated that PilG is recruited to the cell poles (i) by its association with and phosphorylation by ChpA^34^, the Pil-Chp histidine kinase, and (ii) by FimL, a polarly localized scaffold protein^32,34,36,48^. Deletion of either *chpA* or *fimL* decreases PilG polar localization by ∼50%, whereas PilG polar localization is abolished in the *ΔfimL*Δ*chpA* double mutant^34^. These findings suggest that there may be two separate populations of PilG, one associated with ChpA and one associated with FimL. We therefore tested whether ChpA and FimL are required for the localization of ChpB. While *fimL* deletion had no measurable effect on ChpB localization, deletion of *chpA* abolished ChpB polar localization (Fig 4A, B; Suppl Fig 12A, B, E). Thus, our results suggest that only the PilG pool associated with ChpA is required for ChpB polar localization.

**Figure 4.**
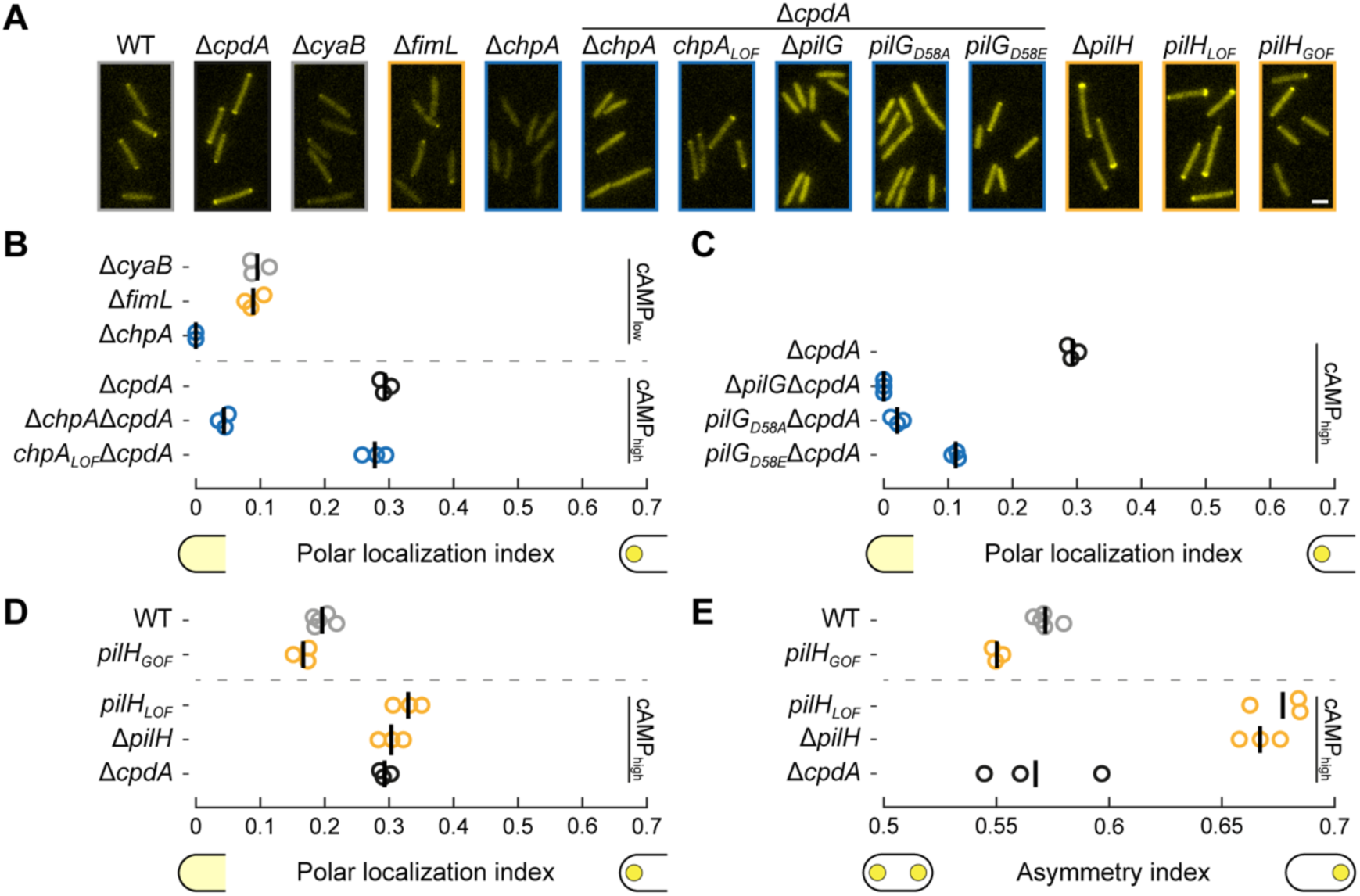
The ChpA histidine kinase and PilH activation affect ChpB localization. **(A)** Representative fluorescence snapshots of motile cells expressing chromosomal ChpB-mNG after 2 h of surface growth. Scale bar, 2 µm. Strains constructed in *ΔcpdA* background are indicated. All other strains are in WT background **(B-C)** Polar localization index of ChpB-mNG in motile cells after 2 h of surface growth in (**B**) *chpA* and *fimL* mutants or in (**C**) *pilG* mutants. *cpdA* was deleted in *chpA* and *pilG* mutants to eliminate potential confounding effects of low cAMP levels. Circles, median of each biological replicate. Vertical bars, mean across biological replicates. **(D-E)** Polar localization index and asymmetry index of motile cells expressing ChpB-mNG in *pilH* mutants after 2 h of surface growth. Circles, median of each biological replicate. Vertical bars, mean across biological replicates. For corresponding fluorescence profiles, asymmetry index and mean cell fluorescence measurements see Suppl Fig 12.

One explanation for the dependence on ChpA is that polar recruitment of ChpB requires phosphorylation of PilG by ChpA. To test this hypothesis, we quantified ChpB localization in a ChpA mutant, *chpA_LOF_*, which is unable to undergo autophosphorylation and subsequent phosphoryl group transfer to PilG^30,34^. Localization of ChpB was indistinguishable from WT in *chpA_LOF_* (Fig 4A, B; Suppl Fig 12A, B, F). These data suggest that phosphorylation of PilG is not required for polar recruitment of ChpB.

We also assessed the role of PilG phosphorylation in ChpB polar recruitment by mutating the PilG phosphoryl group acceptor, aspartate 58, to either alanine or glutamic acid. The *pilG_D58A_ and pilG_D58_*_E_ mutants have decreased twitching motility and cAMP production, suggesting that both point mutations diminish PilG function^34^. When we investigated ChpB localization in *pilG_D58A_ and pilG_D58_*_E_, ChpB polar localization was nearly undetectable in *pilG_D58A_* and greatly decreased in *pilG_D58E_* (Fig 4A, C; Suppl Fig 12A, B, F), even when cAMP levels are restored (Δ*cpdA*). These results further support that ChpB localization depends on PilG because PilG_D58A_ and PilG_D58E_ cannot properly localize to the leading pole^34^. We cannot, however, distinguish whether phosphorylation of PilG contributes to ChpB localization. Overall, we conclude that ChpA and PilG promote localization of ChpB to the leading pole, primarily in a phosphorylation independent manner.

We further investigated the role of PilH phosphorylation in the localization of ChpB-mNG using phosphorylation site mutants of PilH, *pilH_D52A_* and *pilH_D52E_* (hereafter *pilH_LOF_ and pilH_GOF_*). We have previously reported that *pilH_LOF_ and pilH_GOF_* mimic the inactive (non-phosphorylated) and active (phosphorylated) state of PilH, respectively, as assessed by twitching motility, cAMP production, and localization assays^34^. We observed that *pilH_LOF_* phenocopied Δ*pilH* with increased ChpB asymmetric polar localization, while ChpB-mNG localization was comparable to WT in *pilH_GOF_* (Fig 4A, D, E; Suppl Fig 12A, G). As PilH phosphorylation ensures that PilG dynamically localizes to the leading pole, resulting in increased PilG asymmetric localization in *ΔpilH* and *pilH_LOF_*^34^, we infer that PilH∼P indirectly regulates ChpB localization to the leading pole by enabling ChpB dynamic behavior.

### PilG and PilH work together to exclude PilK from the leading pole

To further investigate how the Pil-Chp response regulators segregate PilK to the lagging pole, we monitored the localization of mNG-PilK in phosphorylation mutants of PilG and PilH. Even when cAMP levels were normalized by deleting *cpdA*, PilK polar localization was increased in *pilG_D58A_* and *pilG_D58E_*, with *pilG_D58_*_A_ showing higher PilK polar localization than *pilG_D58_*_E_ (Fig 5A, C; Suppl Fig 12C, D, H). This finding is consistent with both *pilG* mutants having decreased PilG function. In *chpA_LOF_*, a background in which PilG cannot become phosphorylated, polar localization of PilK was also enhanced, comparable to the level of PilK polar localization observed in the *pilG* point mutants (Fig 5A, C; Suppl Fig 12C, D, H). Because polar localization of PilK was higher in Δ*pilG* compared to either of the PilG phosphorylation site mutants (Fig 5A, C), our data suggest that PilG phosphorylation affects the degree of PilK localization to the lagging pole.

**Figure 5.**
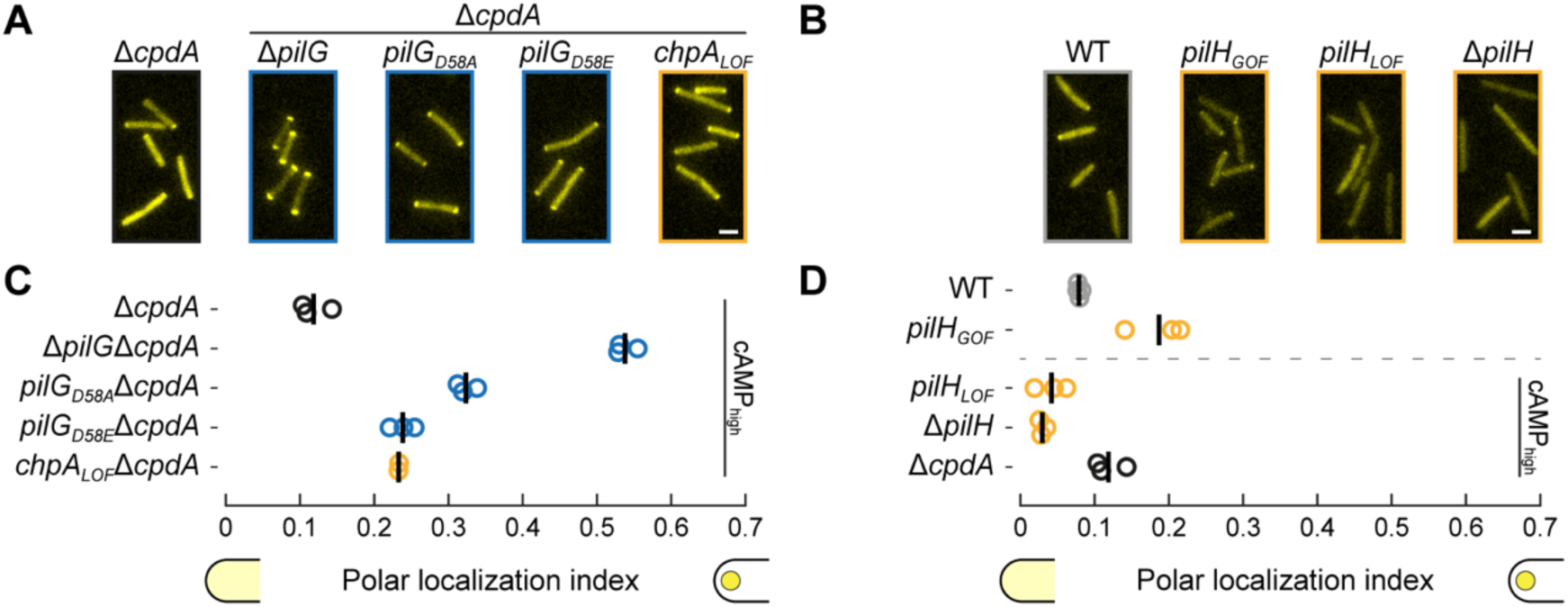
PilG and PilH are required to exclude PilK from the leading pole. **(A-B)** Representative fluorescence snapshots of motile cells with plasmid-expressed mNG-PilK in (**A**) *pilG* mutants and *chpA_LOF_* or in (**B**) *pilH* mutants, all in ΔpilK background, after 2 h of surface growth. Scale bar, 2 µm. **(C-D)** Polar localization index of motile cells expressing mNG-PilK in (**C**) *pilG* mutants and *chpA_LOF_* or in (**D**) *pilH* mutants after 2 h of surface growth. *cpdA* was deleted in *pilG* mutants and *chpA_LOF_* to eliminate potential confounding effects of low cAMP levels. Circles, median of each biological replicate. Vertical bars, mean across biological replicates. For corresponding fluorescence profiles, asymmetry index and mean cell fluorescence measurements see Suppl Fig 12.

For the PilH phosphorylation site mutants, we found that PilK polar localization was nearly abolished in *pilH_LOF_* (Fig 5B, D; Suppl Fig 12C, D, I), similar to the loss of PilK polar localization in Δ*pilH* (Fig 3F). In contrast, PilK polar localization was enhanced in *pilH_GOF_* (Fig 5B, D; Suppl Fig 12C, D, I). PilK was polarly localized in Δ*pilG*, Δ*pilGpilH_LOF_*, and Δ*pilGpilH_GOF_* (Suppl Fig 13A-E), demonstrating that PilH phosphorylation does not affect PilK polar localization in the absence of PilG. In summary, these results highlight that PilG, and indirectly PilH, work together to segregate PilK to the lagging pole in twitching *P. aeruginosa*.

### PilG and PilH regulate PilJ receptor methylation

To link PilJ methylation to the localization patterns of PilK and ChpB established by the Pil-Chp response regulators, we examined the PilJ methylation state in *pilG* and *pilH* mutant strains encoding the functional 3xFlag-PilJ fusion (Suppl Figs 14, 15), using methylation immunoblots (Fig 6 and Suppl Fig 16). In Δ*pilG*, *pilG_D58A_* and *pilG_D58E_*, the fraction of PilJ methylation increased ∼20-30% compared to WT levels (Fig 6A; Suppl Fig 16A). Similar results were observed in a *ΔcpdA* background, indicating that the increased methylation seen in *pilG* mutants is independent of cAMP levels (Fig 6A; Suppl Fig 16A). In the case of Δ*pilG*, the increased fraction of PilJ methylation required PilK and was enhanced in the absence of ChpB (Fig 6C; Suppl Fig 16C). The enhanced fraction of PilJ methylation in the absence of functional PilG is consistent with the role of PilG in recruiting ChpB to the leading pole and segregating PilK to the lagging pole (Figs 3-5).

**Figure 6.**
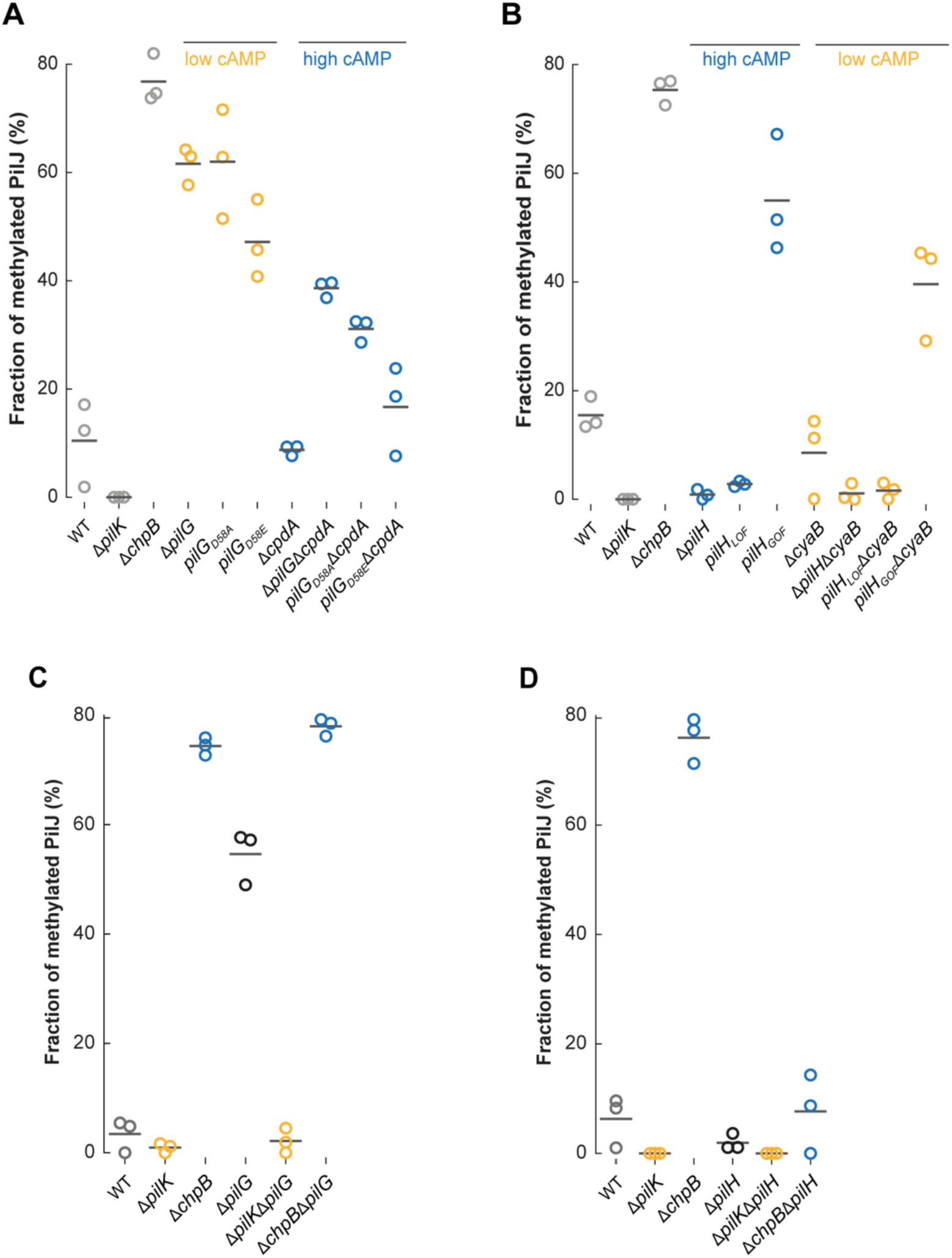
PilG and PilH regulate PilJ receptor methylation. PilJ methylation was quantified as in Fig. 1A after 2 h of surface growth for **(A)** *pilG* mutants **(B)** *pilH* mutants **(C)** *ΔpilGΔpilK* and *ΔpilGΔchpB* double mutants or **(D)** *ΔpilHΔpilK* and *ΔpilHΔchpB* double mutants. **(A-D)** The fraction of methylated PilJ (methylated PilJ signal/total PilJ signal) is represented in the graph and was quantified from 3 biological replicates (circles). Horizontal bar, mean across biological replicates. LOF, loss of function. GOF, gain of function. For corresponding twitching motility assays, cAMP assays, and immunoblot images see Suppl Figs 14-16.

In *ΔpilH* and *pilH_LOF_*, where PilK is cytoplasmic and ChpB asymmetric polar localization is enhanced (Figs 3-5), the fraction of PilJ methylation was decreased, mirroring that of Δ*pilK* and of Δ*pilH*Δ*pilK* (Figs 6B, D; Suppl Fig 16B, D). Compared to *ΔchpB,* PilJ methylation remained low in *ΔpilHΔchpB* (Fig 6D; Suppl Fig 16D), suggesting that cytoplasmic PilK cannot methylate PilJ even in the absence of ChpB. In *pilH_GOF_* (Fig 6B; Suppl Fig 16B), where PilK polar localization was enhanced (Fig 5B, D), PilJ methylation was increased. These data are most consistent with a model in which PilK promotes PilJ methylation at the lagging pole, while ChpB ensures that PilJ demethylation primarily occurs at the leading pole. The spatial control of PilJ methylation would ensure that upon twitching reversals, PilJ at the new leading pole is methylated and “poised” to efficiently promote Pil-Chp signaling once mechanosensing occurs (Figure 7).

**Figure 7.**
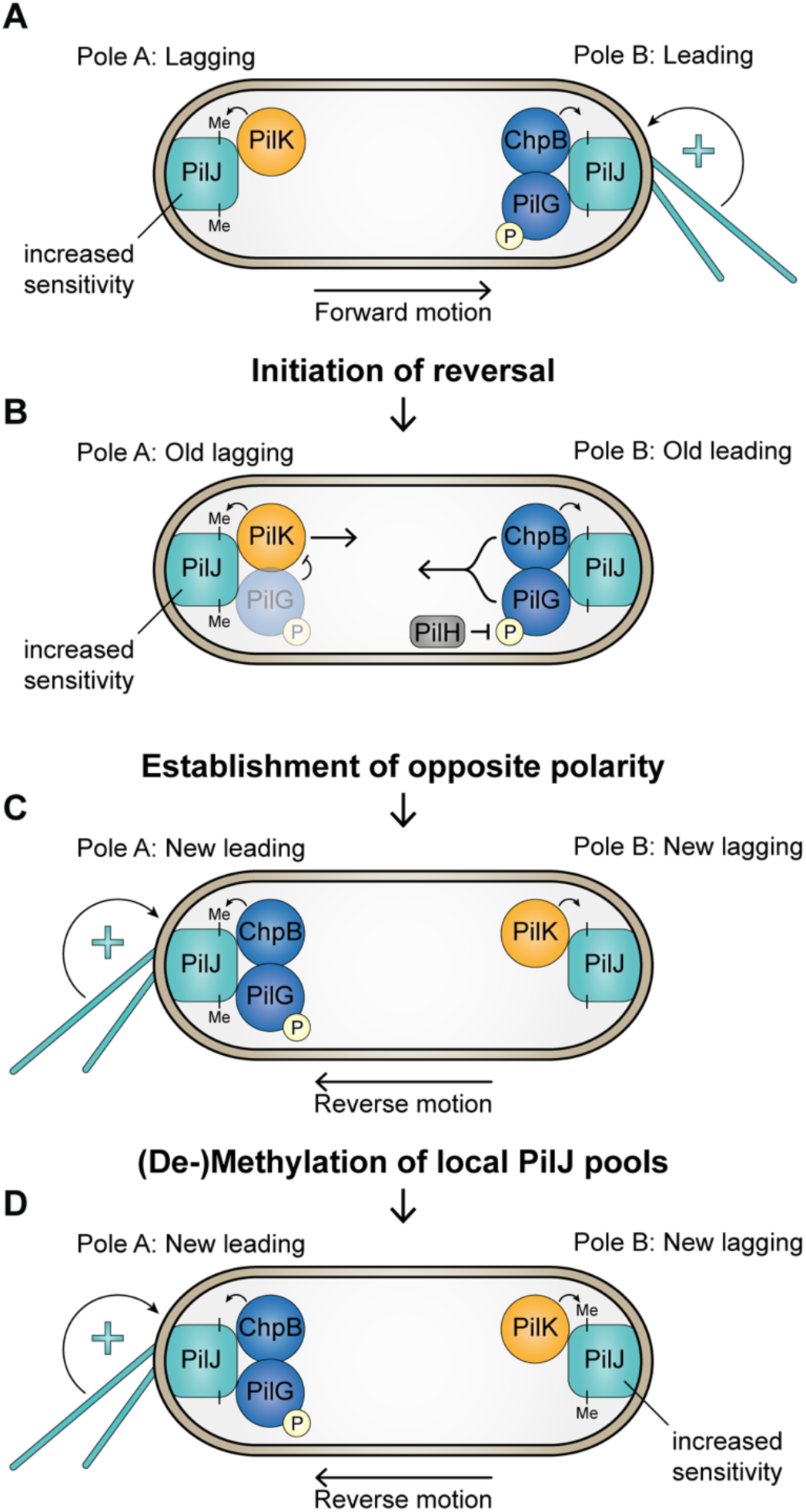
Spatial control of sensory adaptation modulates local signaling hubs of the Pil-Chp system during mechanosensing. **(A)** PilG asymmetric localization establishes the polarity of TFP. TFP are preferentially assembled at the leading pole to propel the cell forward and transmit surface-derived mechanical signals. The mechanical signals, sensed by PilJ, initiate a positive feedback loop that continuously activates the Pil-Chp system at the leading pole^33,34^. The lagging pole has fewer or no TFP and thus less activation of the Pil-Chp system. Leading-pole localized PilG recruits ChpB to the leading pole and excludes PilK from the leading pole, so that PilK is primarily localized to the lagging pole. This subcellular architecture ensures that the pool of PilJ at the leading pole becomes demethylated, while the pool of PilJ at the lagging pole is methylated and poised for activation upon reversal of TFP polarity and twitching motility direction. **(B)** Initiation of a twitching motility reversal. By antagonizing PilG phosphorylation, PilH may trigger re-localization of PilG and ChpB from the leading pole to the lagging pole, thereby initiating a twitching motility reversal. Accumulation of PilG at the old lagging pole excludes PilK from that pole and thereby drives PilK re-localization to the old leading pole. **(C)** Establishment of new leading and lagging poles. Once PilG accumulates at the old lagging pole, PilG establishes a new leading pole, in which methylated PilJ is already primed for efficient activation of the Pil-Chp system. PilG accumulation initiates TFP activity and twitching motility in the opposite direction, completing a twitching motility reversal. ChpB and PilK are now localized to the new leading and lagging pole, respectively. **(D)** Inversion of the local PilJ methylation state. By localizing to the new lagging and leading pole, respectively, PilK and ChpB change the methylation state of the polar pools of PilJ, thereby resetting the sensitivity and activity of PilJ at each pole. This process ensures sensory adaptation to localized mechanical stimuli.

## Discussion

Despite decades of work that have revealed the intricate details of adaptation in bacterial chemotaxis^49^, our understanding of adaptation in other bacterial sensory systems that are widespread throughout nature^50^ is limited. Here, we report the mechanism of sensory adaptation in bacterial mechanosensing, using the Pil-Chp system of *Pseudomonas aeruginosa* as a model system. By controlling the methylation state of the PilJ receptor upon mechanosensing, the PilK methyltransferase and the ChpB methylesterase modulate the two outputs of the Pil-Chp system: twitching motility and cAMP production. Using functional fluorescent fusions, we discovered that PilK and ChpB are spatially segregated to the lagging and leading poles, respectively, in twitching *P. aeruginosa* cells. In addition to the PilK and ChpB catalytic activity, we showed that the *in vivo* PilJ methylation state depends on the localization of both enzymes. We further found that the Pil-Chp response regulators not only regulate TFP polarity and dictate twitching motility reversals, but that PilG, and indirectly PilH, coordinate the spatial segregation of PilK and ChpB. Together, our results reveal how Pil-Chp sensory adaptation uses spatial control as a strategy to modulate mechanosensing in *P. aeruginosa.* Therefore, our work uncovered an unrecognized mechanism to achieve adaptation to sensory stimuli in bacteria.

Despite their small size, *P. aeruginosa* cells can sense spatial environmental information. We and others have revealed that TFP transmit mechanical stimuli derived from surfaces at the leading pole^26,28,33^. The Pil-Chp system, while encoding core components of bacterial sensory pathways, has evolved a complex spatial regulatory network to sense and to respond to polarized mechanical stimuli. Specifically, we demonstrated that PilG and PilH, the antagonistic response regulators of the system, exhibit spatial regulation that is critical for localized Pil-Chp signaling^34^. For example, PilG is dynamically recruited to the leading pole, mirroring the localization of the TFP mechanosensors. This “local” pool of PilG at the leading pole amplifies TFP activity and Pil-Chp signaling, which enables persistent twitching motility and production of cAMP (Fig 7A). Reversals of twitching direction are orchestrated by PilH, the antagonistic Pil-Chp response regulator. Bipolarly localized PilH inhibits the activity of local PilG pools by potentially initiating PilG relocalization to the opposite cell pole. This event triggers a twitching motility reversal and resets localized Pil-Chp signaling (Fig 7B).

Building on this previous work, we now propose that the sensory adaptation branch of the Pil-Chp system also exhibits spatial regulation to facilitate TFP-mediated mechanosensing. Unlike CheR and CheB, which are anchored to MCP arrays at the cell pole independently of the localization of flagella^51,52^, the Pil-Chp sensory adaptation enzymes are segregated to opposite mechanosensory cell poles. PilK localization to the lagging pole ensures that this pool of PilJ is methylated and poised to respond to mechanical stimuli upon TFP polarity and twitching motility reversals. When PilG relocalizes to the lagging pole to establish a new leading pole, the increased sensitivity of methylated PilJ may amplify the signal needed to complete a twitching motility reversal and produce cAMP (Fig 7C). Our model further predicts that the inverse occurs at the opposite pole. Recruitment of ChpB by PilG at the leading pole would dampen PilJ methylation and thus attenuate Pil-Chp signaling (Fig 7D). We propose that Pil-Chp sensory adaptation resets localized Pil-Chp signaling at each pole instead of resetting “global” signaling capabilities of MCPs, as is the case during chemotaxis^49^. Spatial control of PilJ methylation would enable the Pil-Chp system to optimally respond to polarized mechanical stimuli during the establishment of the leading pole. This model would also explain why PilJ methylation modulates twitching motility reversals and the amplitude of cAMP production rather than returning the system to a single baseline.

Beyond their role in the excitatory branch of the Pil-Chp system^26,30–34^, our experiments reveal that the response regulators PilG and PilH have taken the additional function of regulating PilJ methylation and thus sensory adaptation. In the chemotaxis system, CheB phosphorylation by activated CheA enhances CheB methylesterase activity^18^, thereby tilting the “global” balance of MCP methylation to achieve sensory adaptation. In the Pil-Chp system, the ChpB receiver domain lacks key residues involved in phosphotransfer^53^, suggesting that ChpB phosphorylation might not play a role in regulating PilJ methylation. We instead provide evidence that PilJ methylation depends on PilG, and indirectly on PilH, because these CheY-like response regulators function together to segregate the Pil-Chp sensory adaptation enzymes to opposing poles during twitching motility. This model is supported by the observation that in *pilG* and *pilH* mutants, the localization of PilK and ChpB as well as overall PilJ methylation are unregulated. Thus, the mechanisms that ensure interactions between sensory adaptation enzymes and their cognate MCPs likely have profound effects on MCPs methylation and adaptation. Consistent with this idea, CheR and CheB in the *E. coli* chemotaxis system are recruited and anchored to the polar MCP arrays by a C-terminal pentapeptide found in MCPs ^51,52,54,55^. When their cognate C-terminal pentapeptide is deleted, CheR and CheB cannot mediate an adequate sensory adaptation response during chemotaxis^56–58^. As there is no discernible C-terminal pentapeptide in PilJ^59^, we infer that PilK and ChpB gained spatial control of PilJ methylation by tightly linking their localization to response regulators of the system. Our work thus provides a unique perspective on how the excitatory branch of the Pil-Chp system directly regulates sensory adaptation during *P. aeruginosa* mechanosensing. In the chemotaxis system of *Bacillus subtilis*, CheY also mediates adaptation and MCP methylation, but CheY does so indirectly by participating in a feedback loop with additional proteins (CheC and CheD) not found in the *E. coli* chemotaxis and the Pil-Chp systems^60^. By relying on spatial organization and on the response regulators, we suggest that Pil-Chp sensory adaptation has achieved a mechanistic complexity that has primarily been evident in eukaryotic sensory adaptation^3^.

Taken together, our previous and current studies illustrate how *P. aeruginosa* has optimized the excitatory and sensory adaptation branches of the Pil-Chp system to respond to mechanical stimuli, which is required to colonize new habitats, either in its natural environment or in human tissues. The spatial regulatory network that we describe likely ensures that the Pil-Chp system can operate independently from the other three sensory pathways found in *P. aeruginosa*, all of which encode homologous core components^23,61^. In the case of PilK and ChpB, their spatial segregation likely arose during their evolution as they have a unique evolutionary history when compared to their chemotactic counterparts, CheR and CheB^50^. As 90% of bacterial sensory systems encode sensory adaptation enzymes^50^, we anticipate that spatial regulation will be increasingly recognized as a critical component of adaptation in bacterial sensory systems.

## STAR Methods

### Bacterial strains, growth conditions and media

*Pseudomonas aeruginosa* PAO1 ATCC 15692 (American Type Culture Collection) was used for all experiments in this study. *Escherichia coli* strains DH5α and Stellar (Takara Bio) were used for cloning. *E. coli* strain S17.1 was used for conjugative mating with *P. aeruginosa*. All strains were grown in LB medium (Carl Roth) at 37 °C with 280 rpm shaking with the addition of appropriate antibiotics, for *E. coli* 100 µg.ml^-1^ ampicillin/carbenicillin or 10 µg.ml^-1^ gentamycin, for *P. aeruginosa* 300 or 400 µg.ml^-1^ carbenicillin or 60 or 100 µg.ml^-1^ gentamycin. Solid LB media for cell growth, cAMP measurements, and methylation immunoblots were prepared by adding 1.5% (wt · vol^−1^) agar (Fisher Bioreagents) and antibiotics as appropriate. For experiments that included pCuAIgent plasmid-containing strains, no inducers were added to the media.

For subsurface stab assays, 35 ml of sterile 1% LB agar were poured into tissue culture-treated plastic dishes (150 x 25 mm, Corning). The plates were allowed to solidify overnight and used the next day. Before use, plates were dried in a flow hood for 20 min. Semi-solid tryptone media for single-cell twitching and microscopy were prepared by autoclaving 5 g.l^-1^ tryptone (Carl Roth), 2.5 g.l^-1^ NaCl (Fisher Bioreagents), and 0.5 % (wt.vol^−1^) agarose standard (Carl Roth). To reproducibly ensure single-cell twitching, the tryptone media were allowed to cool down to 70 °C in the autoclave followed by cooling to 55 °C for 20 - 30 min in a water bath and pouring 28 ml medium into 90 mm petri dishes. The plates were dried in a flow hood for exactly 30 min and stored for at least one day and no more than two days at 4 °C in a plastic bag.

### Strains and vector construction

Strains used in this study are listed in Key Resources Table 1, plasmids in Key Resources Table 2, and oligonucleotides in Key Resources Table 3. Chromosomal mutants of *P. aeruginosa* were generated as described previously^34,62^. Genes were deleted or integrated by two-step allelic exchange using the suicide vectors pEX18_Amp_ or pEX18_Gent_ following the protocol in ^63^. For genomic in-frame gene deletions or integrations, approximately 500 to 1000 base pair fragments of the up- and downstream regions of the designated gene were combined by PCR amplification and subsequent Gibson assembly^64^. Point mutants were created by integrating the mutated gene into the corresponding gene deletion background by allelic exchange. All mutants, marker-free deletions, and insertions were verified by PCR and sequencing. Fluorescent fusion proteins were generated by fusing mNeonGreen (mNG) to the N- or C-termini of designated genes separated by a 5x-Glycine or 4x-Glycine-1x-Serine linker. Expression and size of fluorescent fusions was verified by Western blot. Functionality was tested by measuring twitching motility in the subsurface stab assay and cAMP production. Plasmids were constructed using standard Gibson assembly^64^ or QuikChange site directed mutagenesis protocols^65^ and introduced into *P. aeruginosa* cells by conjugative mating with *E. coli* S17.1 as donor^63^ or by using electroporation^66^.

mNG-tagged PilK was expressed from a cumic acid-inducible plasmid that was originally designed for *Agrobacterium tumefaciens*^67^. Plasmid pXP313 was used as template for vector backbone PCR amplification and the mNG-PilK fusion fragment amplified from pMK26 was integrated by Gibson assembly to yield plasmid pMK35. The kanamycin resistance gene was exchanged with the gentamycin resistance gene (template pEX18gm) by the same approach, yielding plasmid pMK65 that was used for all plasmid-expressed mNG-PilK experiments. In addition, plasmid pMK66 was generated as an empty vector control. It was modified for easier cloning by inserting a synthetic sequence downstream of the CuO regulatory site. The insert can be excised to linearize the plasmid for Gibson assembly using restriction enzymes XhoI and XbaI.

### Twitching motility subsurface stab assay

Single *P. aeruginosa* colonies were stabbed through 1% LB agar, with appropriate antibiotics if needed, to the bottom of the dish with a sterile 10 µl pipette tip. The plates were incubated at 37 °C overnight, and the agar was carefully removed with a 10 µl inoculating loop. The diameter of the observed twitching motility zone at the bottom of the dish was measured for each biological replicate (in cm). For each biological replicate, the relative twitching motility zone was calculated as a percentage of the WT twitching motility zone.

### cAMP quantification

cAMP levels were measured using the reporter plasmid pUC18-PlacP1/POXB20 as previously described^34^. LacP1 is a synthetic cAMP responsive promoter^31^ and OXB20 is a strong constitutive promoter (Oxford Genetics Ltd. UK, Sigma). *P. aeruginosa* were transformed with the reporter plasmid and plated on LB agar with 400 µg.ml^-1^ Carbenicillin (GoldBio) to generate single colonies. Single colonies were grown overnight in 500 µl of LB broth in a 96-deepwell plate at 37 °C, with shaking at 900 rpm. Overnight cultures were diluted 1:100 and grown for 3 h with shaking, as described above. To measure cAMP levels in mid-log phase liquid conditions, an aliquot of the 1:100 subculture grown for 3 h was fixed by adding methanol-free paraformaldehyde (PFA, Thermo Scientific) to a final concentration of 4% and incubated for 10 min at room temperature. The reaction was quenched by adding 2.5 M glycine (Fisher Scientific) to a final concentration of 0.3 M. Using the same 1:100 subculture, a second fraction of bacteria was spotted onto agar plates and grown for 2 h, or up to 6 h, at 37 °C. Spots were scraped, resuspended in phosphate-buffered saline (PBS; UCSF Media Production Unit) and fixed, as described above. Samples were diluted in PBS and stored at 4 °C for no more than one day before analysis. An LSRFortessa flow cytometer (BD Biosciences) located in the UCSF Parnassus Flow CoLab (RRID:SCR_018206) was used to measure yellow fluorescent protein (YFP) in at least 30,000 mKate2-positive single cells. Exported Flow Cytometry Standard (FCS) files were analyzed in FlowJo and the YFP fluorescence intensity reported as median value for each biological replicate.

### Methylation immunoblots

Single colonies of 3xFlag-PilJ encoding strains were grown overnight in 500 µl of LB broth in a 96-deepwell plate at 37 °C, with shaking at 900 rpm and with appropriate antibiotics as needed. Overnight cultures were diluted 1:100 and grown for 3 h with shaking, as described above. To quantify PilJ methylation during mid-log growth, 500 µl of the 1:100 subculture grown for 3 h was pelleted by centrifugation (10 000 x g, 5 min, 4°C). The pellet was suspended in 2X Laemmli sample buffer (Biorad) with 2.5% 2-mercaptoethanol (Sigma) and 0.1 U/µl Pierce universal nuclease (Thermo Scientific) and frozen at −20 °C. To quantify PilJ methylation during surface growth, a second fraction of bacteria from the same 1:100 subculture grown for 3 h was spotted onto 1.5% agar plates and grown for 2 h at 37 °C. Cells were scraped, resuspended into PBS (UCSF Media Production Unit), collected by centrifugation, suspended in sample buffer, and stored as described above. Pierce™ 660nm Protein Assay Reagent with Ionic Detergent Compatibility Reagent (Thermo Scientific) was used to measure protein concentration of each whole cell lysate. 2.5 µg of each whole cell lysate was separated on 11% SDS PAGE gels containing 0.074% bis-acrylamide (low-bis SDS-PAGE). Gels were electrophoresed at 200 V in running buffer (25 mM Tris, 192 mM glycine, 0.1% SDS, pH 8.3) until the 75 kDa marker of the protein standard (5 µl of Precision Plus Protein™ Standard (Biorad)) was ∼ 2 cm from the bottom of the gel. Gels were transferred (20 V, 7 min) to PVDF membranes using the iBlot™ Dry blotting system (Thermo Scientific). Membranes were blocked in TBS (20mM Tris, 150mM NaCl, pH 7.5)/5% milk overnight. Membranes were incubated with monoclonal mouse anti-Flag M2 antibody (affinity isolated, catalogue number F3165, Sigma) in TBST (20mM Tris, 150mM NaCl, 0.05% Tween20, pH 7.5)/5% milk at a final concentration of ∼1 µg/ml for 1 h. Membranes were washed 3x in TBST and incubated with IRDye® 680RD goat anti-mouse IgG secondary antibody (RRID AB_10956588, catalogue number 926-68070, LI-COR) and imaged on a LI-COR Odyssey CLx system. The signal intensity of individual bands was quantified with ImageStudioLite (LI-COR). The fraction of methylated PilJ was calculated from the signal intensity of the methylated PilJ band (bottom band) divided by the signal intensity of the sum of both PilJ methylated and unmethylated bands 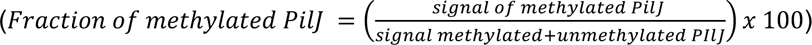

### Immunoblotting

Single colonies of strains expressing mNG protein fusions were grown overnight in LB broth with appropriate antibiotics as needed at 37 °C with shaking. Overnight cultures were diluted 1:100 and grown for 3 h with shaking. 1 ml of the 1:100 subculture was harvested by centrifugation (10 000 x g, 5 min, 4°C). The pellet was suspended in 2X Laemmli sample buffer (Biorad) with 2.5% 2-mercaptoethanol (Sigma) and 0.1 U/µl Pierce universal nuclease (Thermo Scientific) and frozen at −20 °C. Pierce™ 660nm Protein Assay Reagent with Ionic Detergent Compatibility Reagent (Thermo Scientific) was used to measure protein concentration.

2.5 µg of each whole cell lysate and 2 µl of a 1:10 dilution of the Precision Plus Protein™ Standard (Biorad) were separated on Any kD™ Mini-PROTEAN TGX™ Precast SDS PAGE gels. Gels were electrophoresed at 200 V in running buffer (25 mM Tris, 192 mM glycine, 0.1% SDS, pH 8.3) for 40 min. Gels were transferred (20 V, 7 min) to PVDF membranes using the iBlot™ Dry blotting system (Thermo Scientific). Membranes were blocked in in TBS (20mM Tris, 150mM NaCl, pH 7.5)/5% milk overnight, and they were probed with a 1:500 dilution of mouse monoclonal anti-mNG antibody (catalogue number 32F6, Chromotek) in TBST (20mM Tris, 150mM NaCl, 0.05% Tween20, pH 7.5)/5% milk for 1 h. Membranes were washed 3x in TBST and incubated with IRDye® 680RD goat anti-mouse IgG secondary antibody (RRID AB_10956588, catalogue number 926-68070, LI-COR) and imaged on a LI-COR Odyssey system. Immunoblot images were generated using ImageStudioLite (LI-COR).

### Fluorescence microscopy

Microscopy was performed on an inverted Nikon TiE epifluorescence microscope using NISElements (version AR 5.02.03). For phase contrast microscopy, a 40× Plan APO NA 0.9 phase contrast objective was used. For fluorescence microscopy, a 100× Plan APO NA 1.45 phase contrast oil objective and Semrock YFP-2427B filters were used. Microscope settings were identical throughout the study to ensure comparability of fluorescent intensities. Fluorescent images were background subtracted, and snapshots and movies were generated with ImageJ (version 1.53). Data were analyzed with custom scripts using Python (version 3.8.5) and MATLAB (version R2019b) as described previously^34^ and described further below. Custom codes are available on Github (https://github.com/PersatLab/antagonists).

### Cell preparation for single cell twitching experiments

Cells were plated on LB with appropriate antibiotics as needed and incubated overnight at 37 °C. Cells were grown to mid-exponential phase (OD_600_ = 0.2-0.8) in filtered LB medium, with appropriate antibiotics as needed, and diluted to OD_600_ = 0.2. After prewarming the tryptone agarose plates (without antibiotics or inducers for all microscopy experiments) for 45-60 min, a 16 mm round pad was cut out and 1 µl of the diluted cell suspension was pipetted onto the upper side of the agarose pad (the side that was not in contact with the plastic dish bottom). The pads were flipped immediately on a glass bottom dish (P35G-1.5-20-C, MatTek). Four PBS droplets were pipetted to the sides without touching the pad to prevent drying. The samples were used directly for imaging or incubated at 37 °C for imaging within 2 h.

### Quantification of static protein localization

Fluorescent profiles display the mean pixel value of a transversal section of the cell along the mid-cell axis^62^. Cells were prepared and imaged as described above. Several still images of motile cells were recorded to yield hundreds of segmented cells per replicate. Motile cells were typically visible 1 and 2 h after preparation of the sample. Image analysis was performed as described previously^62^ and below. Fluorescent images were acquired at 0, 1 and 2 h after preparation of the microscope dishes. Cells were segmented using phase contrast images with MATLAB-based BacStalk (version 1.8)^68^ and fluorescence profile data was exported as csv files. For comparison of proteins with different expression levels, the profiles were normalized by the total fluorescence of the cell and rescaled to the cell length. Cells were oriented so that the dim pole is at x = −1, the bright pole at x = 1, and mid-cell at x = 0. Mean profiles and standard deviations were computed individually for each biological replicate.

### Polar localization and asymmetry indexes

To quantify the extent of polar localization vs cytoplasmic localization, a polar localization index was derived from the fluorescent profile data. A polar area was defined to measure the polar fluorescence signal 𝐼_A_ and 𝐼_B_ of opposite poles A and B. The polar area was set relative to the ratio between cell width and cell length to account for differences in cell size 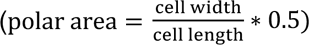. To accurately compute polar localization indexes for proteins with weak polar localization the following correction method was applied. This was necessary because fluorescence profiles of purely cytoplasmic proteins are bell-shaped instead of flat. The fluorescence profile of soluble mNG protein expressed from plasmid pJN105-mNG (uninduced) was used as baseline profile corresponding to a polar localization index of zero^34^. The soluble mNG baseline profile plus standard deviation was subtracted from the measured profiles. The polar localization index is the corrected integrated signal at the defined polar areas *I_AC_* + *I_BC_* divided by the initial total polar fluorescence 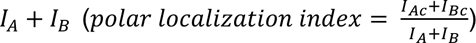. If values slightly below zero occur due to noise, the polar localization index was set to 0. A polar localization index of 0 corresponds to cytoplasmic proteins, and values toward 1 correspond to polarly localized proteins. Values of exactly 1 can never be reached with the applied correction method.

An asymmetry index was determined similarly by taking the ratio between the maximum total fluorescence *I_max_* of opposite poles A and B and the sum of the polar total fluorescence 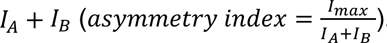. Values around 0.5 correspond to symmetric bipolar localization, a value of 1 corresponds to unipolar localization.

### Reversal frequency of isolated cells and after collisions

Cells were prepared and imaged with phase contrast microscopy as described above. BacStalk was used to segment cells, and image sequences with 5 s interval were recorded for 5 min. The analysis was carried out as described previously^62^. To measure reversal rates, for each frame starting from the first frame in which the cell was classified as moving, the scalar product between the rounded normalized displacement vector 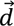 and the cell orientation unit vector 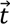 was determined. This yielded a series of numbers corresponding to cellular movement toward the initial leading pole (1), toward the initial lagging pole (−1) or no movement (0) at that timepoint. Timepoints with no movement were removed. Cell movement was considered persistently directional relative to the initial leading pole if the sign remained the same and counted as reversing when a change of sign occurred. To compensate for frequent sign changes occurring for close to non-moving cells, a reversal was only counted when at least two subsequent frames before and after the reversal had the same sign. The displayed reversal frequency is the sum of all considered reversals divided by the total tracked time over all cell tracks for each biological replicate.

Reversals after collisions were counted manually from the raw movies using the ImageJ plugin Cell Counter. A collision was only considered if the cell was moving in the same direction for at least three frames before the collision and the collision lasted for at least two frames (frame interval 5 sec). Collisions with angles below roughly 20° were not considered. Reversals after collision were only considered if the reversal occurred within five frames after the collision ended. Freshly divided cells were not considered. The displayed reversal frequency is the sum of all considered reversals divided by the sum of all considered collisions for each biological replicate.

### Quantification of dynamic protein localization

For dynamic protein localization, image sequences were recorded similarly to still images as described above. Typically, image sequences were recorded after 2 h surface contact with an image interval of 5 s for 2-3 min. Cells were segmented and tracked in a custom code and BacStalk (version 1.8)^68^. Cells were categorized into moving and non-moving subpopulations by applying a speed threshold (here: 26 - 65 nm s^-1^). The positions of both cell poles were determined using the cell outline obtained from BacStalk and poles were labeled according to their position in subsequent frames. Average fluorescence intensities 𝐼_A_ and 𝐼_B_ of opposite poles A and B were measured in an area around the coordinate of each pole with a radius 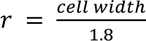. Note, these intensities are not comparable to the polar intensities measured for the static localization because the polar areas are defined differently. Cells were then categorized according to their ratio of average fluorescence intensity between the two poles over the whole cell track 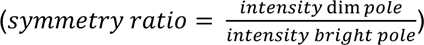. The threshold between symmetric and asymmetric localization was set to 0.8 to allow enough cells in the asymmetric subpopulation for downstream analyses. Cells were considered asymmetric with a ratio below the threshold. From this asymmetric subpopulation an alignment factor was calculated to measure if a cell moves in the direction of the dim or the bright pole. The alignment factor α is the scalar product of a unit vector 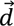 from the dim to the bright pole and the normalized displacement vector 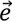. The alignment factor α indicates in which direction the cell is moving relative to the bright pole. An alignment factor α > 0 corresponds to movement toward the bright pole, with α = 1 corresponding to movement exactly parallel to the cell length axis.

### Affinity purification and mass spectrometry

Experiments were performed as previously described^48^. Briefly, LB cultures inoculated from a single colony were grown at 37 °C with vigorous aeration at 250 rpm for 6 h. One hundred microliter of the culture was spread onto 1.5% MinS media^69^ plates and grown overnight at 37 °C. Bacteria were scraped from the plate and resuspended by vortexing and rapid pipetting in 5 mL PBS, and OD_600_ was measured. Cell suspensions were diluted to OD_600_ = 3 in 2 mL PBS, centrifuged at 8000 x g, the supernatants were discarded, and cell pellets were frozen. Pellets were resuspended in 2 ml lysis buffer (50 mM Tris-HCl, pH 7.5, 150 mM NaCl, 0.5% NP-40, 1 mg mL^-1^ lysozyme, protease inhibitor cocktail tablet [Roche], 25 U mL^-1^ benzonase [Invitrogen]). Lysates were incubated on ice for 20 min and sonicated (Branson sonicator 150) with a microtip for 10 s with 1 s manual pulsing (on setting 10, approximately 100 W), three times with minimum 1 min rest intervals on ice. Lysates were centrifuged for 20 min at 14,000 x g, and the soluble portion was decanted and incubated with 50 µl anti-HA beads (as directed by the manufacturer, EZview Red anti-HA affinity gel with antibody clone HA-7, Sigma). Beads were captured by low speed (1000 x g) centrifugation, and the flowthrough was discarded. Bound beads were washed three times with 1 mL of wash buffer (50 mM Tris-HCl pH 7.5, 150 mM NaCl, 0.05% NP-40), vortexed briefly and centrifuged for capture. Beads were washed additionally with buffer lacking NP-40 to remove detergent. Bound material was eluted from beads with 30 µL HA peptide solution (Sigma) in TBS (100 ng/ml). Eluates were flash frozen in liquid nitrogen, thawed, and then then trypsin digested for LC-MS/MS^70^. Digested peptide mixtures were analysed on a Thermo Scientific Velos Pro ion trap MS system equipped with a Proxeon Easy nLC II high-pressure liquid chromatography and autosampler system. Each purification was performed at least four times. AP-MS samples were scored by SAINT^47^.

### Statistical tests and software

To test significance, one-way ANOVA and Tukey’s post hoc tests were done using Python (version 3.8.5). A p-value below 0.05 was considered statistically significant.

## Acknowledgements

The authors would like to thank John S. (Sandy) Parkinson and Caralyn Flack for advice on the methylation immunoblots and Igor Jouline with advice on bioinformatic analysis. J.E., R.P., Y.I., H.M. were supported by funding provided by the NIH (RO1 AI129547, RO1 AI174014, R21 AI154350) and the Cystic Fibrosis Foundation (003224P221, 495008). R.P. was also funded by an NSF GRFP fellowship (1650113) and the NIH (F31 AI147544). I.C. was supported by an NSF GRFP fellowship (2034836). M.K. and A.P. were supported by funding by SNSF 310030-189084. M.K. was also supported by an EMBO postdoctoral fellowship ALTF 495-2020. D.S. was supported by NIH R01AI167412. Any opinions, findings, and conclusions or recommendations expressed in this material are those of the author(s) and do not necessarily reflect the views of the National Science Foundation.

## Author contributions

Conceptualization, R.P., M.K., H.M., Y.I., A.P., J.E.; Data curation, D.S.; Formal analysis, J.vD., J.J., D.S.; Funding acquisition, R.P., M.K., A.P., J.E.; Investigation, R.P., M.K., H.M., Y.I., I.C; Methodology, R.P., M.K., H.M., Y.I., D.S., N.K., A.P., J.E.; Project administration, N.K., A.P., J.E.; Resources, J.vD., J.J., D.S.; Supervision, D.S., N.K., A.P., J.E.; Validation, R.P., M.K., H.M., Y.I., D.S., N.K., A.P., J.E.; Visualization, R.P., M.K., H.M., Y.I., A.P., J.E.; Writing—original draft, R.P., M.K., H.M., Y.I., D.S., A.P., J.E.; Writing-revisions and editing, R.P., M.K., H.M., Y.I., D.S., A.P., J.E.

