## Supplemental figures for "Spatial control of sensory adaptation modulates mechanosensing in *Pseudomonas aeruginosa*"

### Supplemental Information

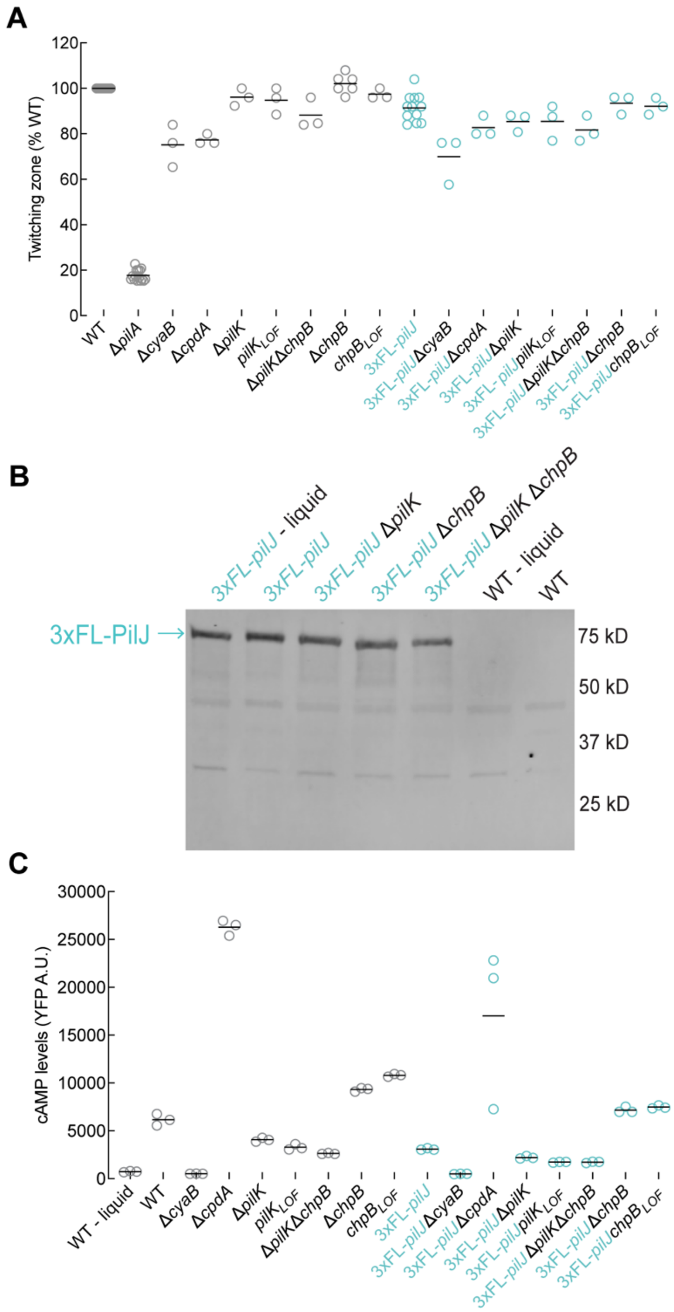


**Supplemental Figure 1. Phenotypic characterization of *3x-Flag-pilJ* fusion strains**
**(A)** Twitching motility of the indicated strains with either the WT *pilJ* or with 3xFL-*pilJ* in the chromosome was quantified by the subsurface stab assay. A representative graph of two independent experiments, with at least three biological replicates per experiment, is shown. Circles, relative twitching motility zone of each biological replicate (% of WT). Horizontal bars, mean across biological replicates. Δ*pilA* serves as a twitching motility deficient control.

**(B)** Immunoblot of whole cell lysates from strains expressing 3xFlag-PilJ (3X-FL-PilJ) from the chromosome separated by conventional SDS-PAGE and immunoblotted with anti-FLAG antibody. Whole cell lysates were prepared from cells grown to mid-log phase in liquid or from cells after 2 h of surface growth. Strains labeled WT have untagged PilJ and serve as a control for the FLAG antibody.

**(C)** cAMP levels of the indicated strains with either the WT *pilJ* or with 3xFL-*pilJ* grown in liquid to mid-log phase or after 2 h of surface growth. cAMP levels were quantified as in Fig 1C. A representative graph of two independent experiments, with three biological replicates per experiment, is shown. Circles, median YFP fluorescence of each biological replicate (~30,000 cells). Horizontal bars, mean across biological replicates. LOF, loss of function.

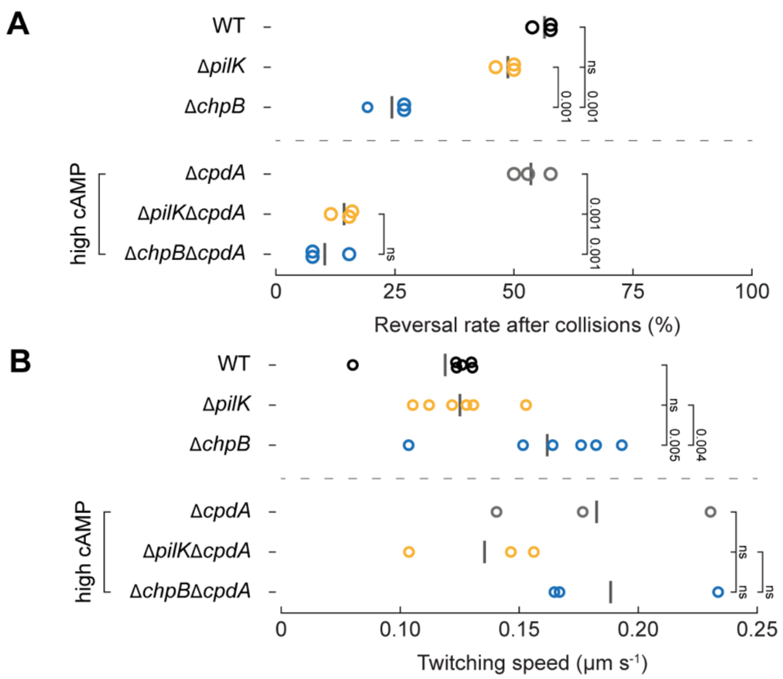


**Supplemental Figure 2. Effect of cAMP levels on single cell twitching reversal rate and single cell twitching speed**

**(A)** Manually counted reversal rates after cell-cell collisions of isolated cells after 2 h surface growth as described in Fig 1B. Circles, median of each biological replicate. Vertical bars, mean across biological replicates.

**(B)** Twitching speed of isolated motile cells was calculated using the cell tracking data from Fig 1B. Circles, median of each biological replicate. Vertical bars, mean across biological replicates. Vertical numbers indicate p-values (ANOVA and Tukey's post hoc test; ns, not significant).

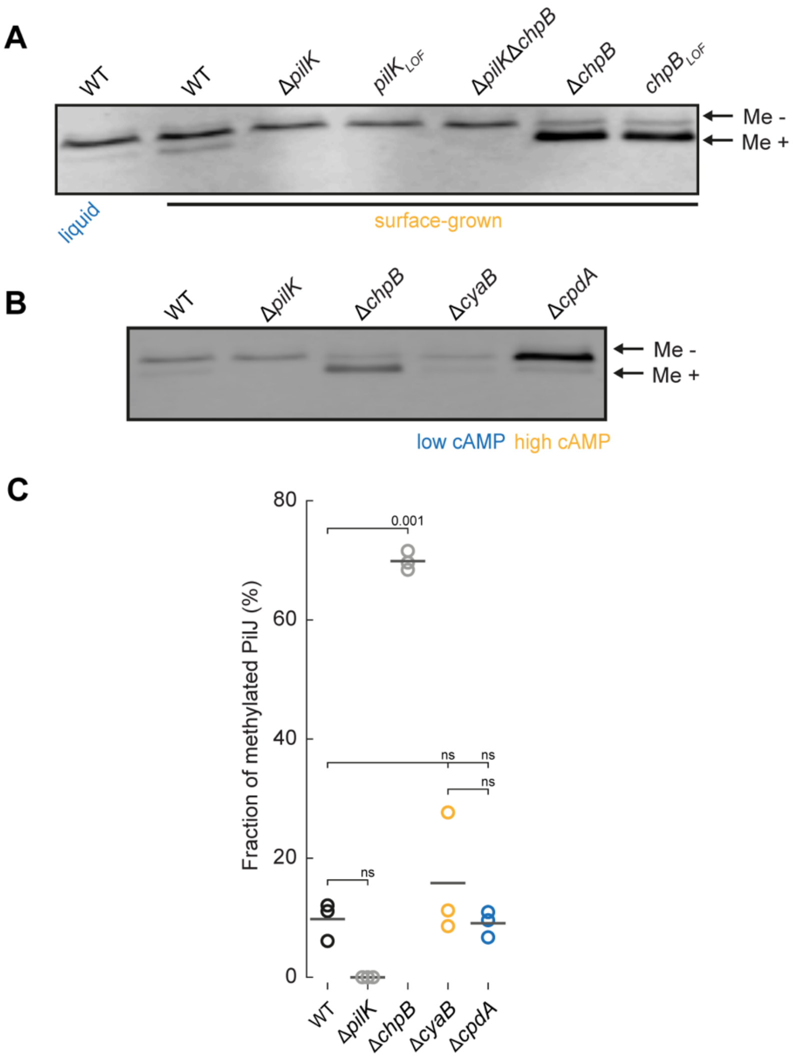


**Supplemental Figure 3. Role of PilK, ChpB, surface exposure, and cAMP production on PilJ methylation.**

**(A)** PilJ methylation increases after 2 h of surface growth and requires PilK and ChpB enzymatic activity. Representative immunoblots of whole cell lysates from strains expressing chromosomal 3xFlag-PilJ separated by "low-bis" SDS-PAGE and immunoblotted with anti-FLAG antibody. Whole cell lysates were prepared from cells grown to mid-log phase in liquid or from cells after 2 h of surface growth. The slower migrating band represents unmethylated PilJ (Me -), while the faster migrating band represents methylated PilJ (Me +). LOF, loss of function.

(**B,C**) Methylation immunoblots (B) and fraction of PilJ methylation (C) is not affected by increased cAMP (Δ*cpdA*) or decreased cAMP (Δ*cyaB*) levels, although the absolute levels of PilJ protein are affected.

The fraction of methylated PilJ (methylated PilJ signal/total PilJ signal) was quantified from 3 biological replicates (circles). Horizontal bar, mean across biological replicates. Numbers indicate p-values (ANOVA and Tukey's post hoc test; ns, not significant).

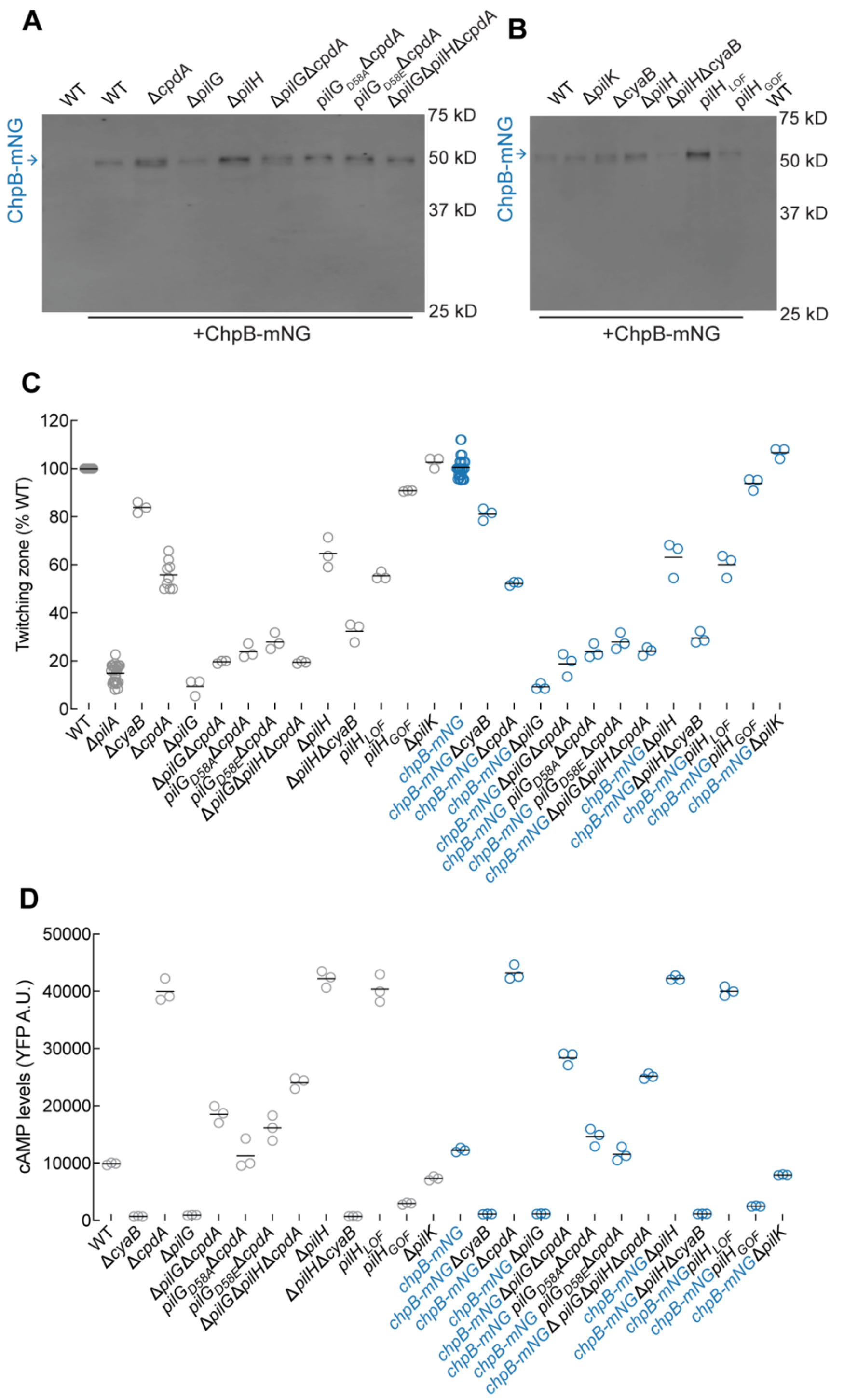


**Supplemental Figure 4. Phenotypic characterization of *chpB-mNG* fusion strains**

**(A)** Immunoblot of whole cell lysates from liquid grown (mid-log phase) strains encoding chromosomally expressed ChpB-mNG and probed with anti-mNG antibody. WT serves as a control for antibody specificity. Steady state levels of ChpB-mNG are increased in strains with elevated cAMP (Δ*pilH*, *pilH*$_{LOF}$, Δ*cpdA*) and decreased in strains with diminished cAMP (Δ*pilG,* Δ*cyaB*).

**(B)** Twitching motility of the indicated strains (WT *chpB* or *chpB-mNG*) was quantified by the subsurface stab assay. A representative graph of two independent experiments, with at least three biological replicates per experiment, is shown. Circles, relative twitching motility zone of each biological replicate (% WT). Horizontal bars, mean across biological replicates. Δ*pilA* serves as a twitching motility deficient control.

**(C)** cAMP levels of the indicated strains (WT *chpB* or *chpB-mNG*) after 2 h surface growth were quantified as in Fig 1C. A representative graph of two independent experiments, with three biological replicates per experiment, is shown. Circles, median YFP fluorescence of each biological replicate (~30,000 cells). Horizontal bars, mean across biological replicates. LOF, loss of function.

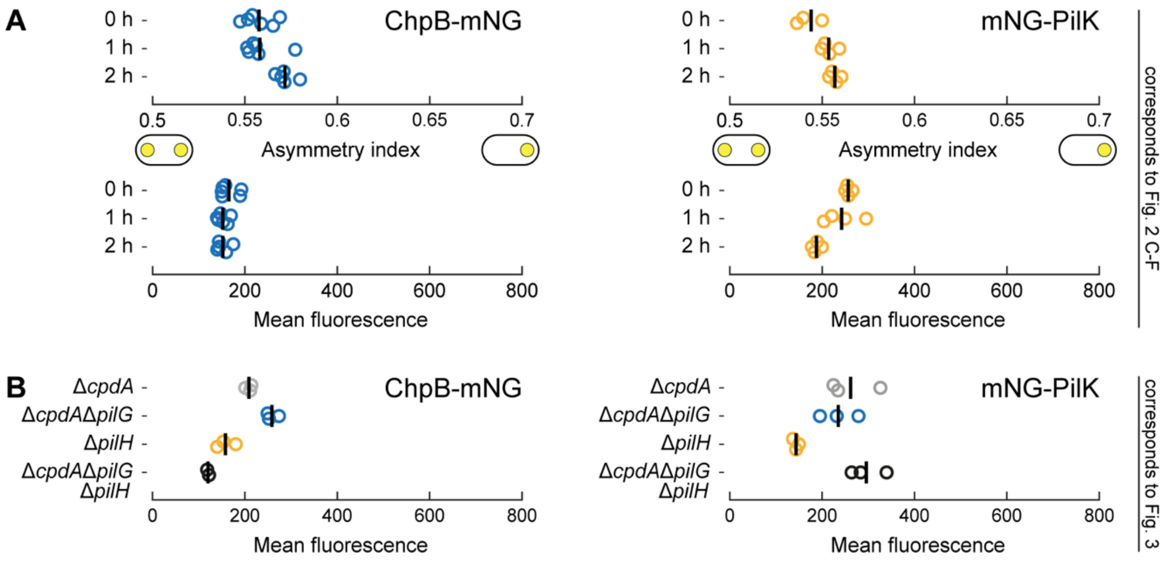


**Supplemental Figure 5. Asymmetry index and fluorescence measurements of PilK and ChpB.**

**(A, B)** Mean cell fluorescence and asymmetry indexes of ChpB-mNG and mNG-PilK. ChpB-mNG is expressed chromosomally from its native locus, while mNG-PilK is expressed from a plasmid. *cpdA* was deleted in *pilG* mutants to eliminate potential confounding effects of low cAMP levels. Panels correspond to main figures as follows: **(A)** Fig. 2 **(B)** Fig. 3. Circles, median of each biological replicate. Vertical bars: mean across biological replicates.

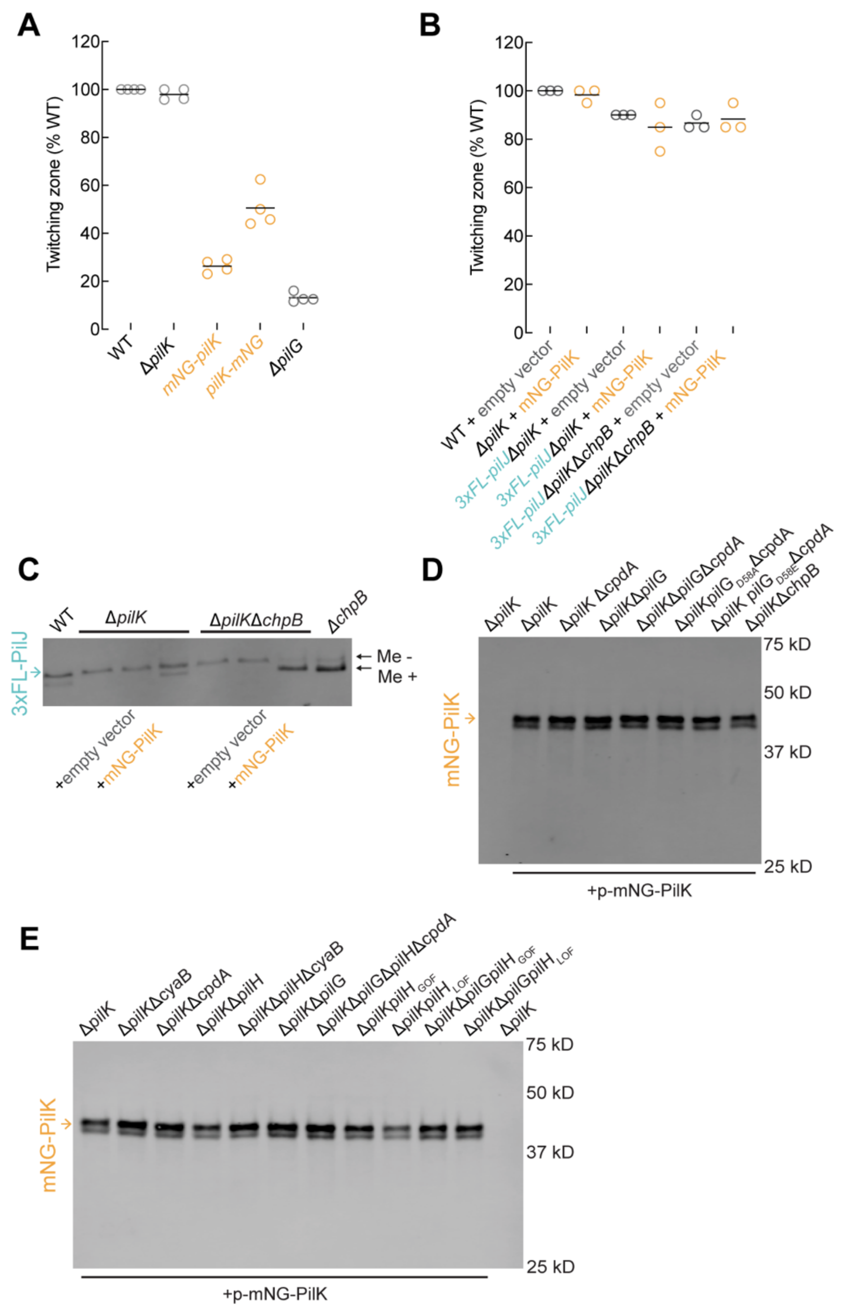


**Supplemental Figure 6. Phenotypic characterization of *mNG-pilK* fusion strains**

**(A-B)** Subsurface stab assays to measure twitching motility in **(A)** strains with chromosomally expressed mNG fusions of PilK and in **(B)** the indicated strains transformed with either empty vector or plasmid-expressed mNG-PilK. A representative graph of two independent experiments, with at least three biological replicates per experiment, is shown. Circles, relative twitching motility zone of each biological replicate (% of WT *P. aeruginosa* PAO1). Horizontal bars, mean across biological replicates. Δ*pilG* is shown in **(A)** as a twitching deficient control.

**(C)** Plasmid-expressed mNG-PilK restores PilJ methylation in *pilK* mutant backgrounds. PilJ methylation was assessed as in Fig 1A in Δ*pilK* and Δ*pilK*Δ*chpB* transformed with either empty vector or with plasmid-expressed mNG-PilK after 2 h of surface growth. WT and Δ*chpB* serve as positive controls for PilJ methylation. The slower migrating band represents unmethylated PilJ (Me-), while the faster migrating band represents methylated PilJ (Me+).

**(D-E)** Immunoblot of whole cell lysates from liquid grown (mid-log phase) Δ*pilK* strains transformed with plasmid-expressed mNG-PilK and probed with anti-mNG antibody. Δ*pilK* in (**D-E)** serves as control for antibody specificity. Expected molecular weight of plasmid expressed mNG-PilK is ~60 kD, while the observed molecular weight is ~40 kD.

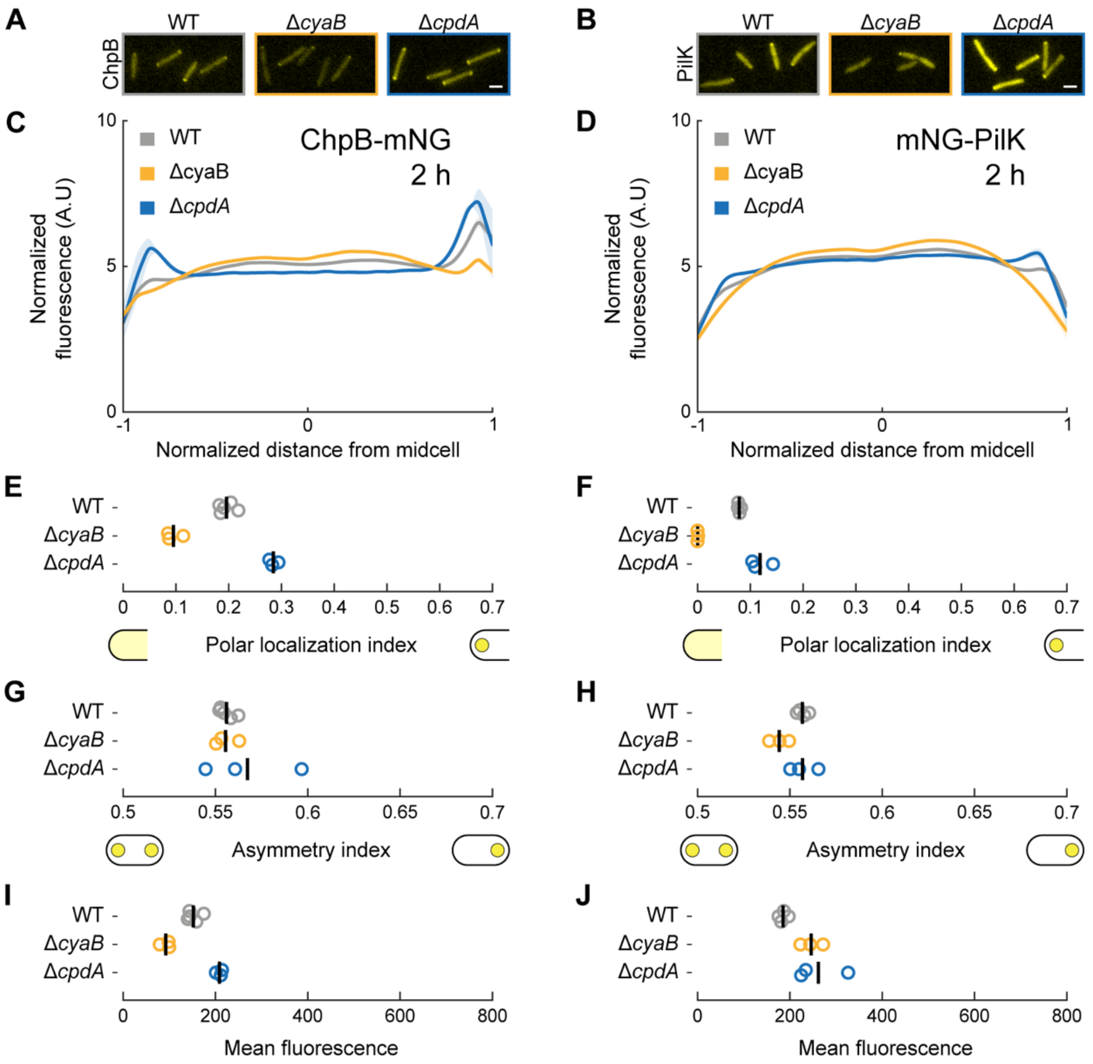


**Supplemental Figure 7. cAMP levels affect the polar localization of ChpB-mNG and mNG-PilK**

**(A, B)** Representative fluorescence snapshots of motile cells after 2 h of surface growth for **(A)** chromosomally expressed ChpB-mNG and **(B)** plasmid-expressed mNG-PilK (in Δ*pilK*). Scale bar, 2 µm.

**(C, D)** Fluorescence profiles of **(C)** ChpB-mNG or **(D)** mNG-PilK in low (Δ*cyaB*) or high (Δ*cpdA*) cAMP backgrounds after 2 h of surface growth. Solid lines, mean normalized fluorescence profiles across biological replicates. Shaded area, standard deviation across biological replicates.

**(G-J)** Corresponding measurements of polar localization index, asymmetry index, and mean cellular fluorescence. Circles, median of each biological replicate. Vertical bars, mean across biological replicates.

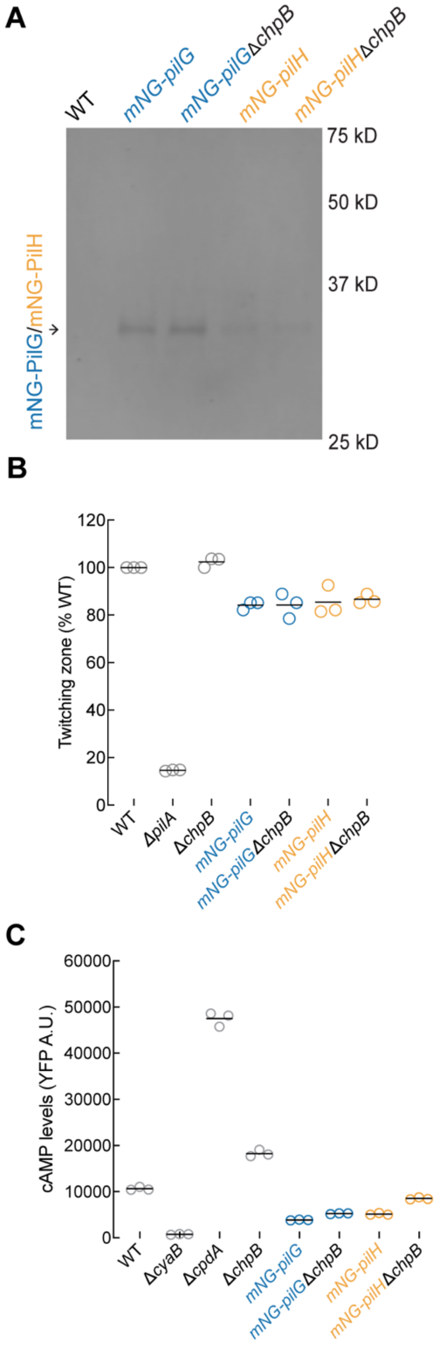


**Supplemental Figure 8. Phenotypic characterization of Δ*chpB* strains expressing *mNG-pilG* or *mNG-pilH***

**(A)** Immunoblot of whole cell lysates from liquid grown (mid-log phase) strains encoding chromosomally expressed mNG-PilG or mNG-PilH and probed with anti-mNG antibody. WT serves as a control for antibody specificity. As mNG-PilG and mNG-PilH have similar molecular weights, arrow points to the band representing either mNG-PilG or mNG-PilH.

**(B)** Subsurface stab assay to measure twitching motility in *mNG-pilG* and *mNG-pilH* fusion strains that lack c*hpB*. A representative graph of two independent experiments, with at least three biological replicates per experiment, is shown. Circles, relative twitching motility zone of each biological replicate (% of WT *P. aeruginosa* PAO1). Horizontal bars, mean across biological replicates. Δ*pilA* serves as a twitching motility deficient control.

**(C)** cAMP levels of *mNG-pilG* and *mNG-pilH* fusion strains in the indicated backgrounds after 2 h of surface growth were quantified as in Fig 1C. A representative graph of two independent experiments, with three biological replicates per experiment, is shown. Circles, median YFP fluorescence of each biological replicate (~30,000 cells). Horizontal bars, mean across biological replicates.

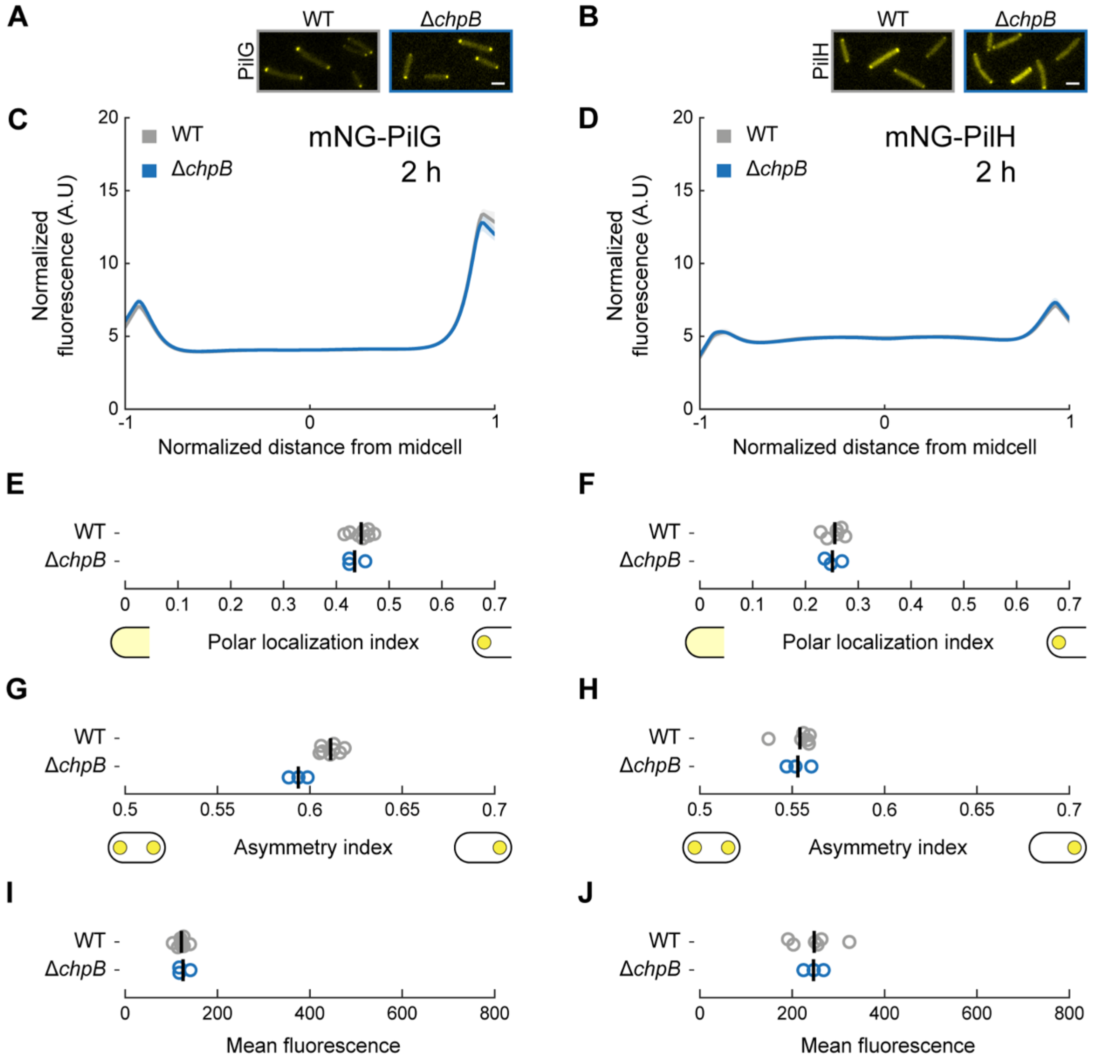


**Supplemental Figure 9. PilG and PilH localization is not affected by ChpB.**

**(A, B)** Representative fluorescence snapshots of motile cells after 2 h of surface growth for **(A)** chromosomally expressed mNG-PilG or **(B)** mNG-PilH. Scale bar, 2 µm.

**(C, D)** Fluorescence profiles of **(C)** mNG-PilG or **(D)** mNG-PilH in Δ*chpB* after 2 h of surface growth. Solid lines, mean normalized fluorescence profiles across biological replicates. Shaded area, standard deviation across biological replicates.

**(E-J)** Corresponding measurements of polar localization index, asymmetry index, and mean cellular fluorescence. Circles, median of each biological replicate. Vertical bars, mean across biological replicates.

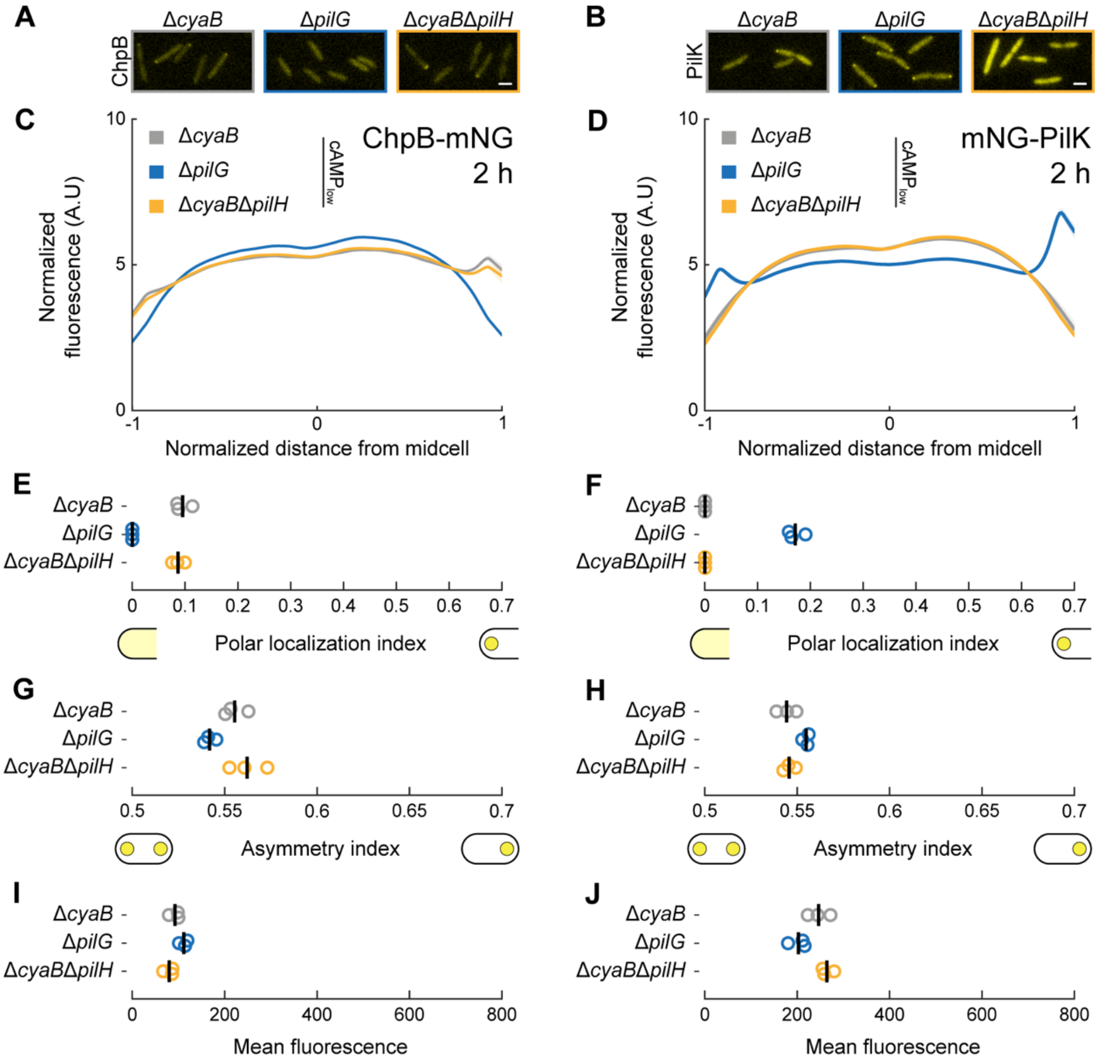


**Supplemental Figure 10. Regulation of ChpB and PilK polar localization by PilG and PilH is not affected by low cAMP levels.**

**(A, B)** Representative fluorescence snapshots of motile cells after 2 h of surface growth for **(A)** chromosomally expressed ChpB-mNG or **(B)** plasmid-expressed mNG-PilK in Δ*pilK*. Scale bar, 2 μm.

**(C, D)** Fluorescence profiles of **(C)** ChpB-mNG or **(D)**mNG-PilK in Δ*pilG* and Δ*pilH* mutants with low cAMP levels (Δ*cyaB* where required) after 2 h of surface growth. Solid lines, mean normalized fluorescence profiles across biological replicates. Shaded area, standard deviation across biological replicates.

**(E-J)** Corresponding measurements of polar localization index, asymmetry index, and mean cellular fluorescence. Circles, median of each biological replicate. Vertical bars, mean across biological replicates.

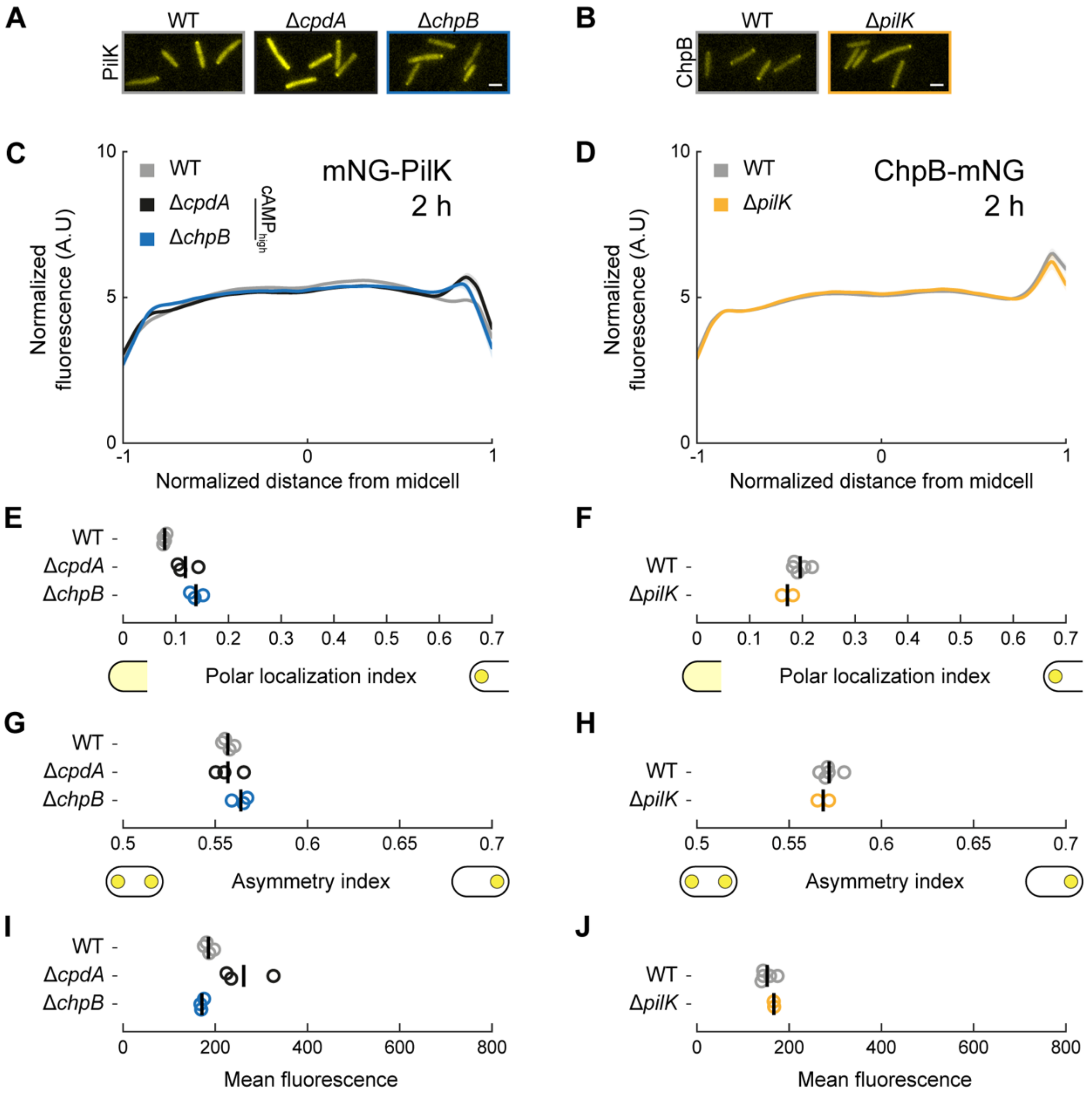


**Supplemental Figure 11. PilK and ChpB localization are independent of each other.**

**(A, B)** Representative fluorescence snapshots of motile cells after 2 h of surface growth for **(A)** plasmid-expressed mNG-PilK in Δ*pilK* or **(B)** chromosomally expressed ChpB-mNG. Scale bar, 2 µm.

**(C, D)** Fluorescence profiles of **(C)** mNG-PilK in Δ*chpB* or **(D)** chromosomally expressed ChpB-mNG in Δ*pilK* after 2 h of surface growth. WT and Δ*cpdA* are shown as reference in **(A, C)** because cAMP levels are increased in Δ*chpB* (cf. Suppl. Fig. 1C). Solid lines, mean normalized fluorescence profiles across biological replicates. Shaded area, standard deviation across biological replicates.

**(E-J)** Corresponding measurements of polar localization index, asymmetry index and mean cellular fluorescence. Circles, median of each biological replicate. Vertical bars, mean across biological replicates.

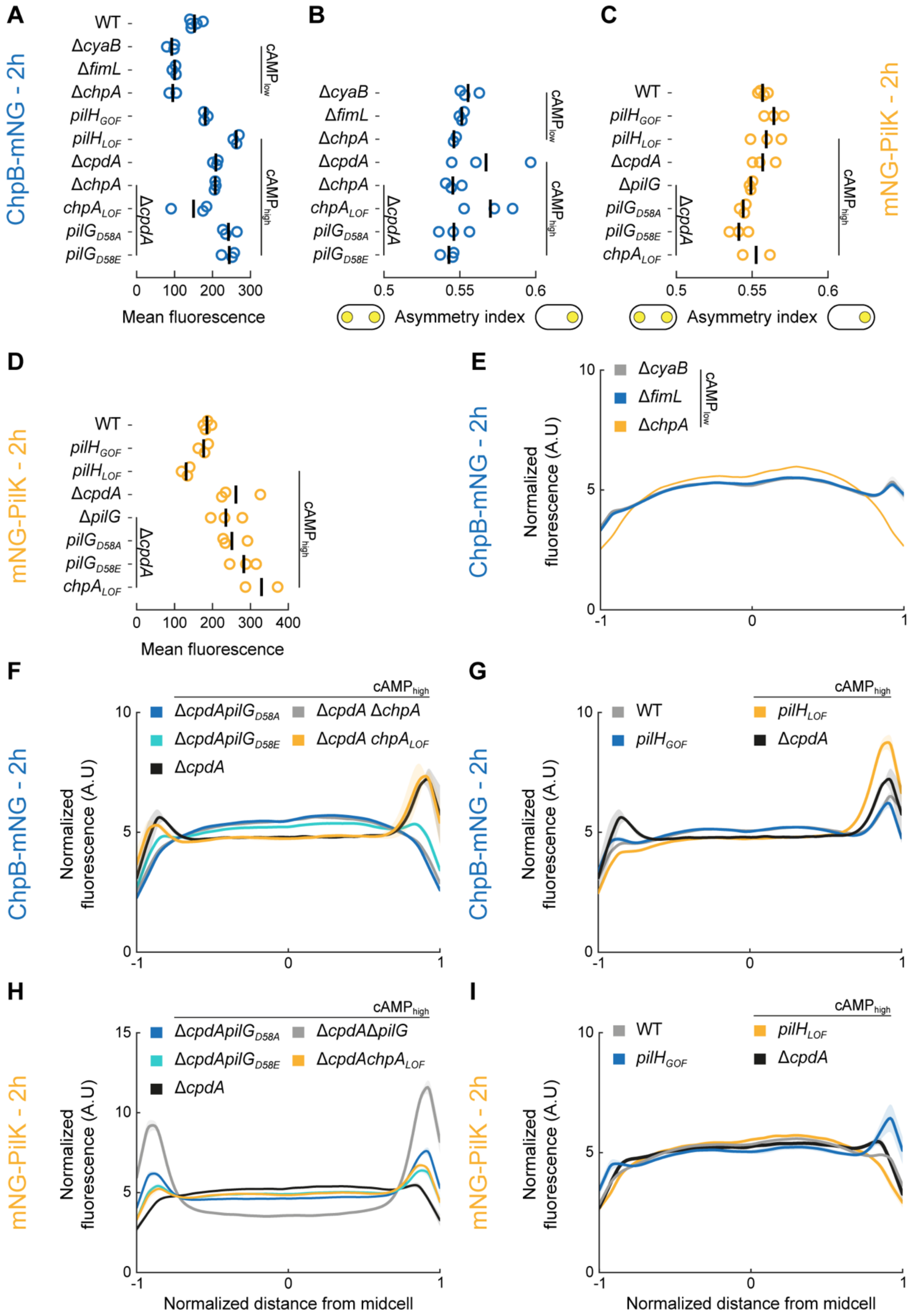


**Supplemental Figure 12. Fluorescence profiles, mean cell fluorescence and asymmetry indices of ChpB-mNG and mNG-PilK in indicated mutant backgrounds**

**(A-D)** Measurements of mean cellular fluorescence **(A,D)** and asymmetry index **(B,C)** for **(A,B)** chromosomally expressed ChpB-mNG or **(C,D)** plasmid-expressed mNG-PilK (in Δ*pilK*) after 2 h of surface growth, corresponding to data shown in Figs 4 and 5. Circles, median of each biological replicate. Vertical bars, mean across biological replicates.

**(E-I)** Fluorescence profiles of **(E,F,G)** chromosomally expressed ChpB-mNG or **(H,I)** plasmid-expressed mNG-PilK after 2 h of surface growth, corresponding to data shown in Figs 4 and 5. Solid lines, mean normalized fluorescence profiles across biological replicates. Shaded area, standard deviation across biological replicates. LOF, loss of function. GOF, gain of function.

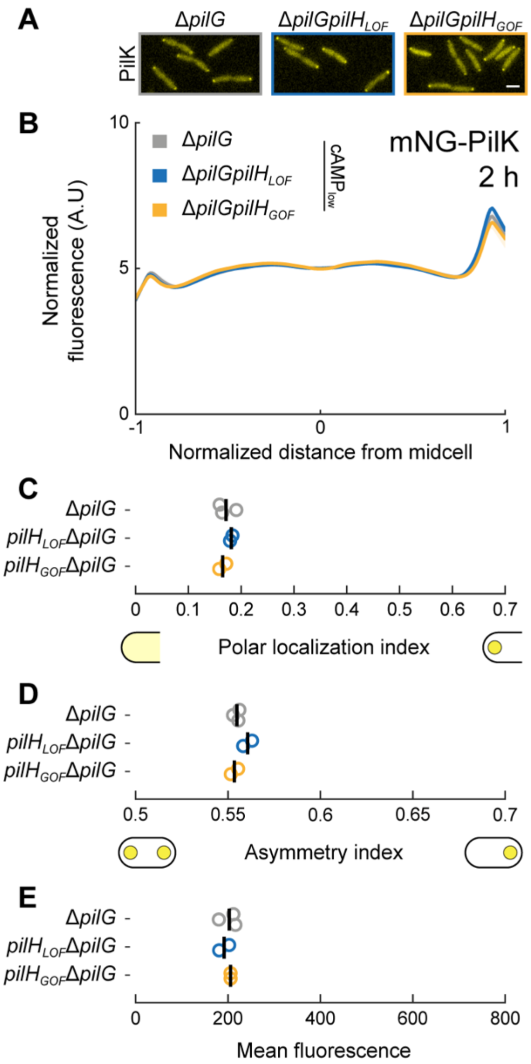


**Supplemental Figure 13. PilH acts on PilG to regulate PilK polar localization.**
**(A)** Representative fluorescence snapshots of cells after 2 h of surface growth for plasmid-expressed mNG-PilK in Δ*pilK*. Scale bar, 2 µm.
**(B)** Fluorescence profiles of plasmid-expressed mNG-PilK in Δ*pilG, pilH*$_{LOF}$Δ*pilG,* or *pilH*$_{GOF}$ Δ*pilG* after 2 h of surface growth. All indicated strains also lack *pilK*. Solid lines, mean normalized fluorescence profiles across biological replicates. Shaded area, standard deviation across biological replicates. LOF, loss of function. GOF, gain of function.
**(C-E)** Corresponding measurements of polar localization index, asymmetry index, and mean cellular fluorescence. Circles, median of each biological replicate. Vertical bars, mean across biological replicates.

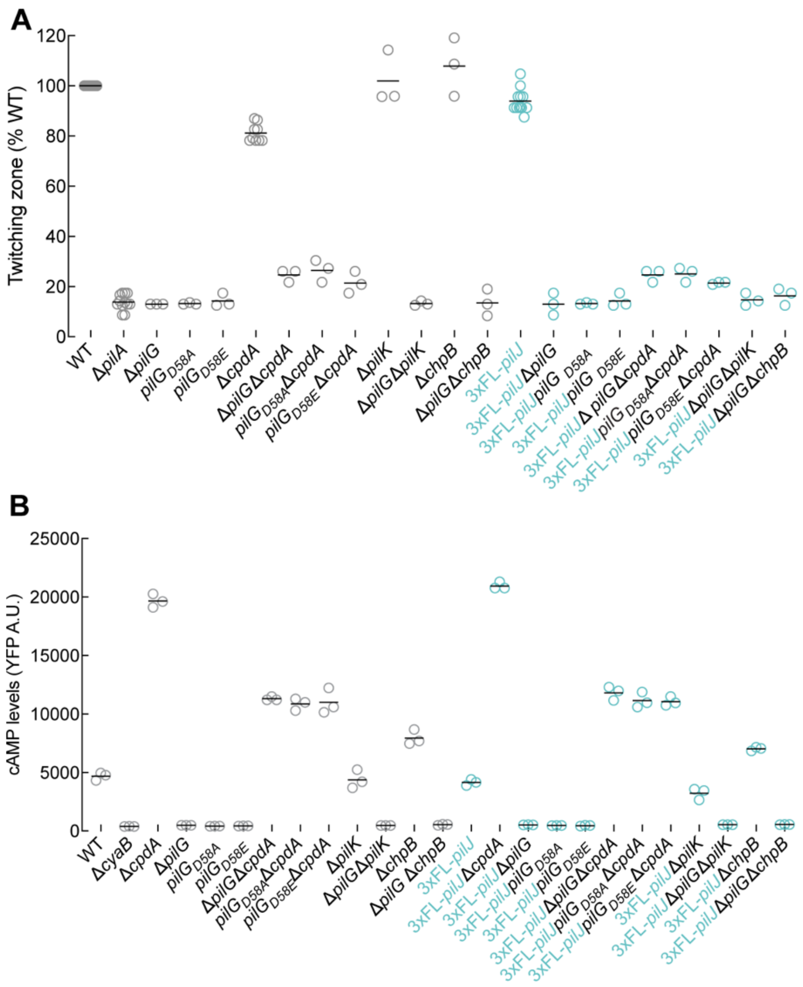


**Supplemental Figure 14. Phenotypic characterization of *3x-Flag-pilJ* fusion strains carrying the indicated *pilG* mutations**

**(A)** Twitching motility of the indicated *pilG* mutants with either the WT *pilJ* or with 3xFL-*pilJ* in the native chromosomal locus was quantified by the subsurface stab assay. A representative graph of two independent experiments, with at least three biological replicates per experiment, is shown. Circles, relative twitching motility zone of each biological replicate (% of WT). Horizontal bars, mean across biological replicates. Δ*pilA* serves as a twitching motility deficient control.

**(B)** cAMP levels of the indicated *pilG* mutants with either the WT *pilJ* or with 3xFL-*pilJ* in the chromosome after 2 h of surface growth were quantified as in Fig 1C. A representative graph of two independent experiments, with three biological replicates per experiment, is shown. Circles, median YFP fluorescence of each biological replicate (~30,000 cells). Horizontal bars, mean across biological replicates.

LOF, loss of function. GOF, gain of function.

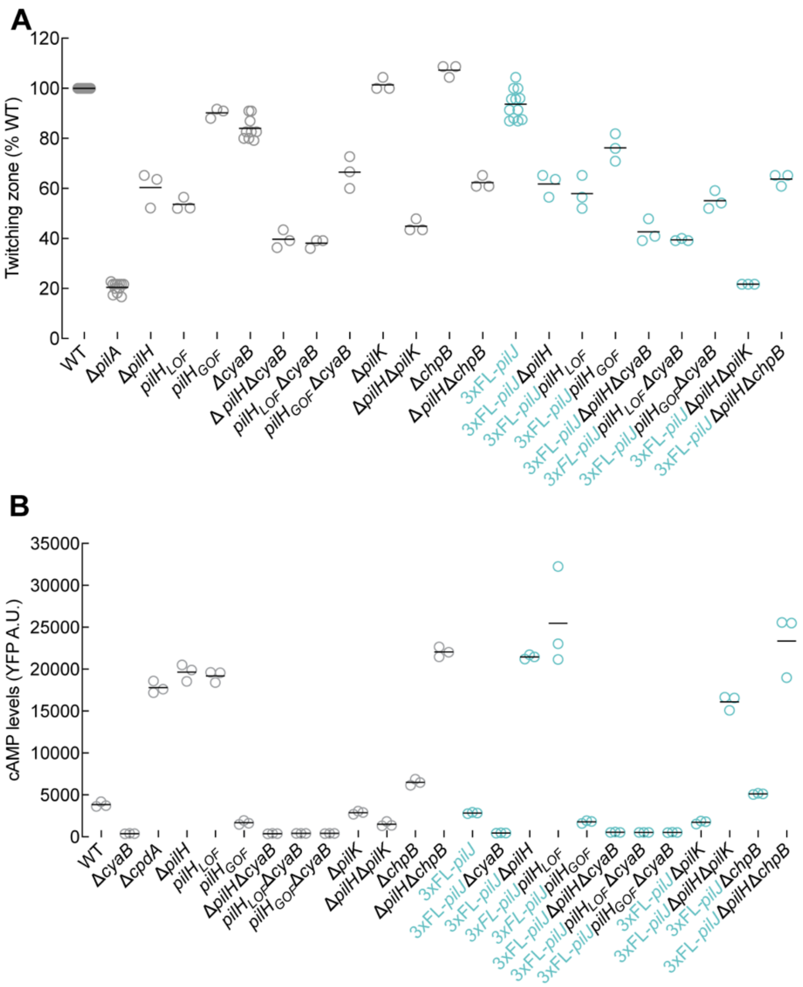


**Supplemental Figure 15. Phenotypic characterization of *3x-Flag-pilJ* fusion strains carrying the indicated *pilH* mutations**

**(A)** Twitching motility of the indicated *pilH* mutants with either the WT *pilJ* or with 3xFL-*pilJ* in the native chromosomal locus was quantified by the subsurface stab assay. A representative graph of two independent experiments, with at least three biological replicates per experiment, is shown. Circles, relative twitching motility zone of each biological replicate (% of WT). Horizontal bars, mean across biological replicates. Δ*pilA* serves as a twitching motility deficient control.

**(B)** cAMP levels of the indicated *pilH* mutants with either the WT *pilJ* or with 3xFL-*pilJ* in the chromosome after 2 h of surface growth were quantified as in Fig 1C. A representative graph of two independent experiments, with three biological replicates per experiment, is shown. Circles, median YFP fluorescence of each biological replicate (~30,000 cells). Horizontal bars, mean across biological replicates. LOF, loss of function. GOF, gain of function.

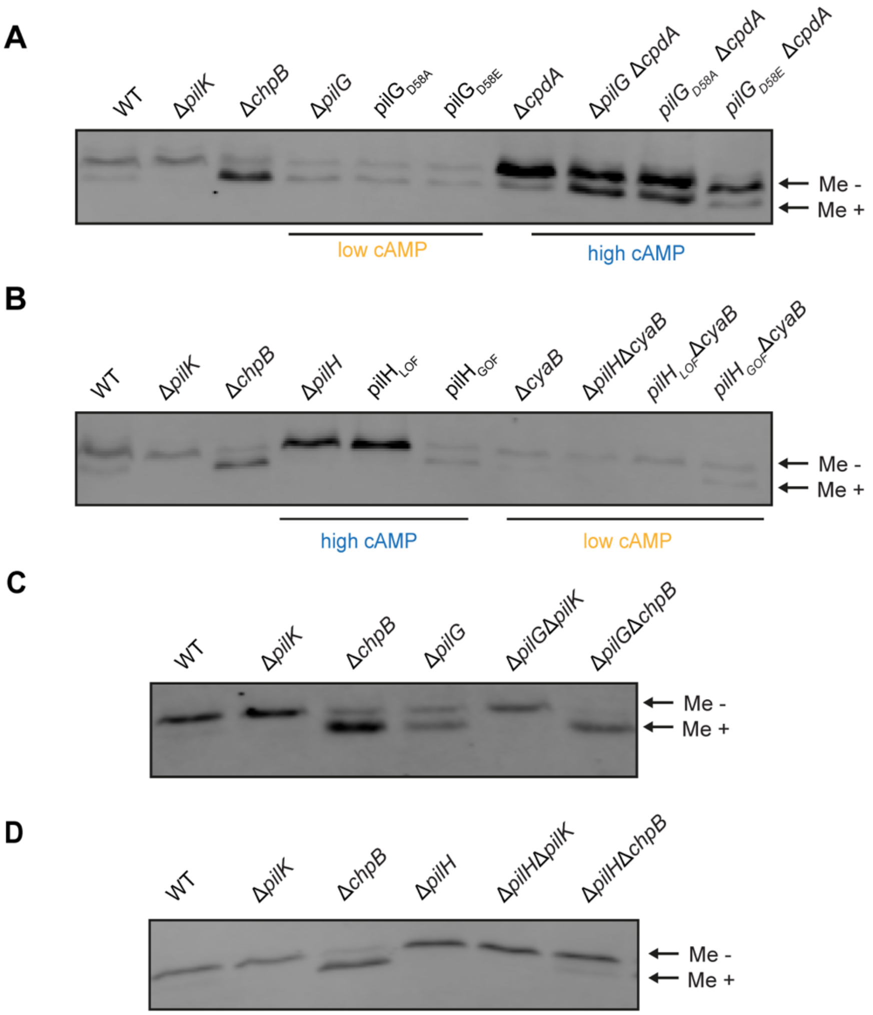


**Supplemental Figure 16. PilG and PilH regulate PilJ receptor methylation.**

**(A-D)** Representative immunoblots of whole cell lysates from strains expressing chromosomal 3xFL-PilJ separated by “low-bis” SDS-PAGE and immunoblotted with anti-FLAG antibody. Whole cell lysates were prepared from cells grown after 2 h of surface growth. The slower migrating band represents unmethylated PilJ (Me -), while the faster migrating band represents methylated PilJ (Me +). PilJ methylation was assessed for **(A)** *pilG* mutants, for **(B)** *pilH* mutants, for **(C)** Δ*pilG*Δ*pilK* and Δ*pilG*Δ*chpB* double mutants, or for **(D)** Δ*pilH*Δ*pilK* and Δ*pilH*Δ*chpB* double mutants. LOF, loss of function. GOF, gain of function.

**Supplemental Table 1. AP-MS reveals a high confidence interaction between PilG and ChpB**

| Bait | Prey | Counts | Replicates | Control Counts | SAINT | BFDR |
|---|---|---|---|---|---|---|
| PilG | PilG | 373 | 3 | 0 | 1 | 0 |
| PilG | FimV | 81 | 3 | 0 | 1 | 0 |
| PilG | FimL | 28.33 | 3 | 0 | 1 | 0 |
| PilG | ChpB | 2.67 | 3 | 0 | 0.98 | 0.01 |
| PilH | PilH | 329 | 3 | 1 | 1 | 0 |
| PilH | FimV | 0 | 3 | 0 | NA | NA |
| PilH | FimL | 0 | 3 | 0 | NA | NA |
| PilH | ChpB | 0 | 3 | 0 | NA | NA |

Lysates from strains expressing overexpressed PilG-HA or PilH-HA were affinity purified and subjected to MS-MS. Shown are preys of interest for PilG-HA and PilH-HA. Previously characterized PilG interactions by AP-MS (PilG:FimV, PilG:FimL)[48] from the same experiment were included in the SAINT scoring[47] and are shown for comparison. SAINT scores closer to 1 and BFDR (Bayesian False Discover Rate) ≤ 0.05 suggest a high confidence interaction.
