## Supplemental tables for "Spatial control of sensory adaptation modulates mechanosensing in *Pseudomonas aeruginosa*"

### Key Resources Tables

#### Key Resources Table 1: Strains used in this study.

| **Name and relevant genotype** | **Source / Reference** | **Identifier** |
|---|---|---|
| Pseudomonas aeruginosa PAO1 | Holloway & Morgan, 1986[71] | RP86<br>MK914 (internal)<br>ATCC 15692 |
| PAO1 pUC18_PlacP1-YFP/POXB20-mKate2 | Kühn *et al*, 2023[34] | HM413 |
| Escherichia coli DH5α (hsdR rec lacZYA ϕ80 lacZM15) | Invitrogen | Na |
| Escherichia coli strain S17.1 (thi pro hsdR recA RP4-2(Tc::Mu)(Km::Tn7)) | Stratagene | Na |
| PAO1 Δ*fliC* (in-frame deletion of PA1092) | Bertrand *et al*, 2010[30] | MK177 |
| PAO1 Δ*pilK* (in-frame deletion of PA0412) | this study | RP151<br>MK571 |
| PAO1 Δ*pilK* pUC18_PlacP1-YFP/POXB20-mKate2 | this study | RP325 |
| PAO1 $pilK_{LOF}$ (PA0412 with substituted residue R92E) | this study | RP393 |
| PAO1 $pilK_{LOF}$ pUC18_PlacP1-YFP/POXB20-mKate2 | this study | RP411 |
| PAO1 Δ*chpB* (in-frame deletion of PA0414) | this study | RP156<br>MK570 |
| PAO1 Δ*chpB* pUC18_PlacP1-YFP/POXB20-mKate2 | this study | RP326 |
| PAO1 $chpB_{LOF}$ (PA0414 with substituted residue H184Y) | this study | RP387 |
| PAO1 $chpB_{LOF}$ pUC18_PlacP1-YFP/POXB20-mKate2 | this study | RP413 |
| PAO1 Δ*pilK* Δ*chpB* | this study | RP189 |
| PAO1 Δ*pilK* Δ*chpB* pUC18_PlacP1-YFP/POXB20-mKate2 | this study | RP327 |
| PAO1 Δ*pilG* (in-frame deletion of PA0408) | Bertrand *et al*, 2010[30] | MK226 |
| PAO1 Δ*pilG* pUC18_PlacP1-YFP/POXB20-mKate2 | Kühn *et al*, 2023[34] | HM421 |
| PAO1 Δ*pilG* Δ*cpdA* | this study | RP565 |
| PAO1 Δ*pilG* Δ*cpdA* pUC18_PlacP1-YFP/POXB20-mKate2 | this study | RP597 |
| PAO1 $pilG_{D58A}$ (loss-of-function mutation D58A) | this study | RP721 |

| | | |
|---|---|---|
| PAO1 *pilG*$_{D58A}$ pUC18_PlacP1-YFP/POXB20-mKate2 | this study | RP779 |
| PAO1 *pilG*$_{D58A}$ Δ*cpdA* | this study | RP761 |
| PAO1 *pilG*$_{D58A}$ Δ*cpdA* pUC18_PlacP1-YFP/POXB20-mKate2 | this study | RP797 |
| PAO1 *pilG*$_{D58E}$ (attempted gain-of-function mutation D58E, behaves like loss-of-function) | this study | RP723 |
| PAO1 *pilG*$_{D58E}$ pUC18_PlacP1-YFP/POXB20-mKate2 | this study | RP780 |
| PAO1 *pilG*$_{D58E}$ Δ*cpdA* | this study | RP763 |
| PAO1 *pilG*$_{D58E}$ Δ*cpdA* pUC18_PlacP1-YFP/POXB20-mKate2 | this study | RP798 |
| PAO1 Δ*pilG* Δ*pilK* | this study | RP360 |
| PAO1 Δ*pilG* Δ*pilK* pUC18_PlacP1-YFP/POXB20-mKate2 | this study | RP403 |
| PAO1 Δ*pilG* Δ*chpB* | this study | RP362 |
| PAO1 Δ*pilG* Δ*chpB* pUC18_PlacP1-YFP/POXB20-mKate2 | this study | RP404 |
| PAO1 Δ*pilH* (in-frame deletion of PA0409) | Barken *et al*, 2008[72] | MK178 |
| PAO1 Δ*pilH* pUC18_PlacP1-YFP/POXB20-mKate2 | Kühn *et al*, 2023[34] | HM422 |
| PAO1 Δ*pilH* Δ*cyaB* | this study | RP567 |
| PAO1 Δ*pilH* Δ*cyaB* pUC18_PlacP1-YFP/POXB20-mKate2 | this study | RP592 |
| PAO1 *pilH*$_{D52A}$ (loss-of-function mutation D52A) | this study | RP734 |
| PAO1 *pilH*$_{D52A}$ pUC18_PlacP1-YFP/POXB20-mKate2 | this study | RP786 |
| PAO1 *pilH*$_{D52A}$ Δ*cyaB* | this study | RP809 |
| PAO1 *pilH*$_{D52A}$ Δ*cyaB* pUC18_PlacP1-YFP/POXB20-mKate2 | this study | RP815 |
| PAO1 *pilH*$_{D52E}$ (gain-of-function mutation D52E) | this study | RP725 |
| PAO1 *pilH*$_{D52E}$ pUC18_PlacP1-YFP/POXB20-mKate2 | this study | RP781 |
| PAO1 *pilH*$_{D52E}$ Δ*cyaB* | this study | RP755 |
| PAO1 *pilH*$_{D52E}$ Δ*cyaB* pUC18_PlacP1-YFP/POXB20-mKate2 | this study | RP794 |
| PAO1 Δ*pilH* Δ*pilK* | this study | RP561 |
| PAO1 Δ*pilH* Δ*chpB* | this study | RP385 |

| | | |
|---|---|---|
| PAO1 Δ*chpA* (in-frame deletion PA0413) | Holloway & Morgan, 1986[71] | MK170 |
| PAO1 Δ*cyaB* (in-frame deletion of PA3217) | Inclan *et al*, 2011[36] | MK174 |
| PAO1 Δ*cyaB* pUC18_PlacP1-YFP/POXB20-mKate2 | Kühn *et al*, 2023[34] | HM416 |
| PAO1 Δ*cpdA* (in-frame deletion of PA4969) | Inclan *et al*, 2011[36] | MK180 |
| PAO1 Δ*cpdA* pUC18_PlacP1-YFP/POXB20-mKate2 | Kühn *et al*, 2023[34] | HM419 |
| PAO1 Δ*fimL* (in-frame deletion of PA1822) | Whitchurch *et al*, 2005[73] | MK179 |
| PAO1 Δ*pilA* (in-frame deletion of PA1822) | Bertrand *et al*, 2010[30] | RP173<br>MK81 |
| PAO1 Δ*fliC* Δ*pilK* | this study | MK1143 |
| PAO1 Δ*fliC* Δ*chpB* | this study | MK1144 |
| PAO1 Δ*fliC* Δ*cpdA* | Kühn *et al*, 2021[33] | MK337 |
| PAO1 Δ*fliC* Δ*pilK* Δ*cpdA* | this study | MK1185 |
| PAO1 Δ*fliC* Δ*chpB* Δ*cpdA* | this study | MK1186 |
| PAO1 *chpB-mNG* (C-terminal fluorescent fusion to mNeonGreen, GGGGG linker, native locus) | this study | RP354<br>MK838 |
| PAO1 *chpB-mNG* pUC18_PlacP1-YFP/POXB20-mKate2 | this study | RP400 |
| PAO1 *pilK-mNG* (C-terminal fluorescent fusion to mNeonGreen, GGGGG linker, native locus) | this study | MK836 |
| PAO1 *mNG-pilK* (N-terminal fluorescent fusion to mNeonGreen, GGGGG linker, native locus) | this study | MK835 |
| PAO1 Δ*fliC* *mNG-pilG* (N-terminal fluorescent fusion to mNeonGreen, GGGGG linker, native locus) | Kühn *et al*, 2021[33] | MK923 |
| PAO1 *mNG-pilG* | Kühn *et al*, 2023[34] | HM240 |
| PAO1 *mNG-pilG* pUC18_PlacP1-YFP/POXB20-mKate2 | Kühn *et al*, 2023[34] | HM441 |
| PAO1 *mNG-pilG* Δ*chpB* | this study | RP193<br>MK1719 |
| PAO1 *mNG-pilG* Δ*chpB* pUC18_PlacP1-YFP/POXB20-mKate2 | this study | RP776 |
| PAO1 Δ*fliC* *mNG-pilH* (N-terminal fluorescent fusion to mNeonGreen, GGGGG linker, native locus) | Kühn *et al*, 2021[33] | MK315 |

| | | |
|---|---|---|
| PAO1 *mNG-pilH* | Kühn *et al*, 2023[34] | HM58 |
| PAO1 *mNG-pilH* pUC18_PlacP1-YFP/POXB20-mKate2 | Kühn *et al*, 2023[34] | HM436 |
| PAO1 *mNG-pilH* Δ*chpB* | this study | RP198<br>MK1720 |
| PAO1 *mNG-pilH* Δ*chpB* pUC18_PlacP1-YFP/POXB20-mKate2 | this study | RP777 |
| PAO1 Δ*fliC* *chpB-mNG* (C-terminal fluorescent fusion to mNeonGreen, GGGGG linker, native locus) | this study | MK1153 |
| PAO1 Δ*fliC* *chpB-mNG* Δ*cyaB* | this study | MK1324 |
| PAO1 Δ*fliC* *chpB-mNG* Δ*cyaB* pUC18_PlacP1-YFP/POXB20-mKate2 | this study | RP889 |
| PAO1 Δ*fliC* *chpB-mNG* Δ*cpdA* | this study | MK1325 |
| PAO1 Δ*fliC* *chpB-mNG* Δ*cpdA* pUC18_PlacP1-YFP/POXB20-mKate2 | this study | RP890 |
| PAO1 Δ*fliC* *chpB-mNG* Δ*pilK* | this study | MK1354 |
| PAO1 *chpB-mNG* Δ*pilK* | this study | RP429 |
| PAO1 *chpB-mNG* Δ*pilK* pUC18_PlacP1-YFP/POXB20-mKate2 | this study | RP440 |
| PAO1 Δ*fliC* *chpB-mNG* Δ*pilG* | this study | MK1286 |
| PAO1 *chpB-mNG* Δ*pilG* | this study | RP425 |
| PAO1 *chpB-mNG* Δ*pilG* pUC18_PlacP1-YFP/POXB20-mKate2 | this study | RP438 |
| PAO1 Δ*fliC* *chpB-mNG* Δ*pilG* Δ*cpdA* | this study | MK1294 |
| PAO1 Δ*fliC* *chpB-mNG* Δ*pilG* Δ*cpdA* pUC18_PlacP1-YFP/POXB20-mKate2 | this study | RP891 |
| PAO1 Δ*fliC* *chpB-mNG* Δ*pilH* | this study | MK1287 |
| PAO1 *chpB-mNG* Δ*pilH* | this study | RP740 |
| PAO1 *chpB-mNG* Δ*pilH* pUC18_PlacP1-YFP/POXB20-mKate2 | this study | RP789 |
| PAO1 Δ*fliC* *chpB-mNG* Δ*pilH* Δ*cyaB* | this study | MK1295 |
| PAO1 Δ*fliC* *chpB-mNG* Δ*pilH* Δ*cyaB* pUC18_PlacP1-YFP/POXB20-mKate2 | this study | RP892 |
| PAO1 Δ*fliC* *chpB-mNG* Δ*pilG* Δ*pilH* Δ*cpdA* | this study | MK1841 |

| | | |
|---|---|---|
| PAO1 Δ*fliC chpB-mNG* Δ*pilG* Δ*pilH* Δ*cpdA* pUC18_PlacP1-YFP/POXB20-mKate2 | this study | RP876 |
| PAO1 Δ*fliC chpB-mNG* Δ*cpdA* Δ*chpA* | this study | MK1738 |
| PAO1 Δ*fliC chpB-mNG* Δ*cpdA* Δ*chpA* pUC18_PlacP1-YFP/POXB20-mKate2 | this study | RP879 |
| PAO1 Δ*fliC chpB-mNG* Δ*cpdA* $chpA_{LOF}$ (loss-of-function mutations D2086A, D2087A, G2088A) | this study | MK1764 |
| PAO1 Δ*fliC chpB-mNG* Δ*cpdA* $chpA_{LOF}$ pUC18_PlacP1-YFP/POXB20-mKate2 | this study | RP877 |
| PAO1 *chpB-mNG* $pilH_{LOF}$ | this study | RP743<br>MK1712 |
| PAO1 *chpB-mNG* $pilH_{LOF}$ pUC18_PlacP1-YFP/POXB20-mKate2 | this study | RP790 |
| PAO1 *chpB-mNG* $pilH_{GOF}$ | this study | MK1713 |
| PAO1 *chpB-mNG* $pilH_{GOF}$ pUC18_PlacP1-YFP/POXB20-mKate2 | this study | RP785 |
| PAO1 *chpB-mNG* Δ*cpdA* $pilG_{D58A}$ | this study | MK1708 |
| PAO1 *chpB-mNG* Δ*cpdA* $pilG_{D58A}$ pUC18_PlacP1-YFP/POXB20-mKate2 | this study | RP791 |
| PAO1 *chpB-mNG* Δ*cpdA* $pilG_{D58E}$ | this study | MK1711 |
| PAO1 *chpB-mNG* Δ*cpdA* $pilG_{D58E}$ pUC18_PlacP1-YFP/POXB20-mKate2 | this study | RP792 |
| PAO1 Δ*fliC pCuAIgent-empty* (cumic acid-inducible plasmid, empty insert, $Gm^R$, pMK66) | this study | MK1770 |
| PAO1 Δ*fliC* Δ*pilK pCuAIgent-mNG-pilK* (N-terminal fluorescent fusion to mNeonGreen, GGGGS linker, expression from cumic acid-inducible plasmid, used without induction, $Gm^R$, pMK65) | this study | MK1659 |
| PAO1 Δ*fliC* Δ*pilK pCuAIgent-mNG-pilK* Δ*cyaB* | this study | MK1744 |
| PAO1 Δ*fliC* Δ*pilK pCuAIgent-mNG-pilK* Δ*cpdA* | this study | MK1745 |
| PAO1 Δ*fliC* Δ*pilK pCuAIgent-mNG-pilK* Δ*pilG* | this study | MK1763 |
| PAO1 Δ*fliC* Δ*pilK pCuAIgent-mNG-pilK* Δ*pilG* Δ*cpdA* | this study | MK1660 |
| PAO1 Δ*fliC* Δ*pilK pCuAIgent-mNG-pilK* Δ*pilH* | this study | MK1661 |
| PAO1 Δ*fliC* Δ*pilK pCuAIgent-mNG-pilK* Δ*pilH* Δ*cpdA* | this study | MK1662 |

| | | |
|---|---|---|
| PAO1 Δ*fliC* Δ*pilK pCuAIgent-mNG-pilK* Δ*pilG* Δ*pilH* Δ*cpdA* | this study | MK1864 |
| PAO1 Δ*fliC* Δ*pilK pCuAIgent-mNG-pilK* Δ*chpB* | this study | MK1746 |
| PAO1 Δ*fliC* Δ*pilK pCuAIgent-mNG-pilK* $pilH_{LOF}$ (loss-of-function mutation D52A) | this study | MK1863 |
| PAO1 Δ*fliC* Δ*pilK pCuAIgent-mNG-pilK* $pilH_{LOF}$ (gain-of-function mutation D52E) | this study | MK1862 |
| PAO1 Δ*fliC* Δ*pilK pCuAIgent-mNG-pilK* Δ*cpdA* $pilG_{D58A}$ (loss-of-function mutation D58A) | this study | MK1865 |
| PAO1 Δ*fliC* Δ*pilK pCuAIgent-mNG-pilK* Δ*cpdA* $pilG_{D58E}$ (attempted gain-of-function mutation D58E, behaves like loss-of-function) | this study | MK1866 |
| PAO1 Δ*fliC* Δ*pilK pCuAIgent-mNG-pilK* $pilH_{LOF}$ Δ*pilG* | this study | MK1910 |
| PAO1 Δ*fliC* Δ*pilK pCuAIgent-mNG-pilK* $pilH_{LOF}$ Δ*pilG* | this study | MK1909 |
| PAO1 3xFLAG-*pilJ* (PA0411 N-terminus fused with 3xFLAG tag, GGGGG linker) | this study | RP491 |
| PAO1 3xFLAG-*pilJ* pUC18_PlacP1-YFP/POXB20-mKate2 | this study | RP494 |
| PAO1 3xFLAG-*pilJ* Δ*pilK* | this study | RP495 |
| PAO1 3xFLAG-*pilJ* Δ*pilK* pUC18_PlacP1-YFP/POXB20-mKate2 | this study | RP501 |
| PAO1 3xFLAG-*pilJ* Δ*chpB* | this study | RP497 |
| PAO1 3xFLAG-*pilJ* Δ*chpB* pUC18_PlacP1-YFP/POXB20-mKate2 | this study | RP502 |
| PAO1 3xFLAG-*pilJ* Δ*pilK* Δ*chpB* | this study | RP499 |
| PAO1 3xFLAG-*pilJ* Δ*pilK* Δ*chpB* pUC18_PlacP1-YFP/POXB20-mKate2 | this study | RP503 |
| PAO1 3xFLAG-*pilJ* $pilK_{LOF}$ | this study | RP516 |
| PAO1 3xFLAG-*pilJ* $pilK_{LOF}$ pUC18_PlacP1-YFP/POXB20-mKate2 | this study | RP530 |
| PAO1 3xFLAG-*pilJ* $chpB_{LOF}$ | this study | RP518 |
| PAO1 3xFLAG-*pilJ* $chpB_{LOF}$ pUC18_PlacP1-YFP/POXB20-mKate2 | this study | RP531 |
| PAO1 3xFLAG-*pilJ* Δ*cyaB* | this study | RP539 |
| PAO1 3xFLAG-*pilJ* Δ*cyaB* pUC18_PlacP1-YFP/POXB20-mKate2 | this study | RP581 |

| | | |
|---|---|---|
| PAO1 3xFLAG-*pilJ* Δ*cpdA* | this study | RP547 |
| PAO1 3xFLAG-*pilJ* Δ*cpdA* pUC18_PlacP1-YFP/POXB20-mKate2 | this study | RP585 |
| PAO1 3xFLAG-*pilJ* Δ*pilG* | this study | RP512 |
| PAO1 3xFLAG-*pilJ* Δ*pilG* pUC18_PlacP1-YFP/POXB20-mKate2 | this study | RP528 |
| PAO1 3xFLAG-*pilJ* Δ*pilG* Δ*cpdA* | this study | RP596 |
| PAO1 3xFLAG-*pilJ* Δ*pilG* Δ*cpdA* pUC18_PlacP1-YFP/POXB20-mKate2 | this study | RP598 |
| PAO1 3xFLAG-*pilJ* $pilG_{D58A}$ | this study | RP727 |
| PAO1 3xFLAG-*pilJ* $pilG_{D58A}$ pUC18_PlacP1-YFP/POXB20-mKate2 | this study | RP782 |
| PAO1 3xFLAG-*pilJ* $pilG_{D58A}$ Δ*cpdA* | this study | RP759 |
| PAO1 3xFLAG-*pilJ* $pilG_{D58A}$ Δ*cpdA* pUC18_PlacP1-YFP/POXB20-mKate2 | this study | RP796 |
| PAO1 3xFLAG-*pilJ* $pilG_{D58E}$ | this study | RP729 |
| PAO1 3xFLAG-*pilJ* $pilG_{D58E}$ pUC18_PlacP1-YFP/POXB20-mKate2 | this study | RP783 |
| PAO1 3xFLAG-*pilJ* $pilG_{D58E}$ Δ*cpdA* | this study | RP764 |
| PAO1 3xFLAG-*pilJ* $pilG_{D58E}$ Δ*cpdA* pUC18_PlacP1-YFP/POXB20-mKate2 | this study | RP799 |
| PAO1 3xFLAG-*pilJ* Δ*pilG* Δ*pilK* | this study | RP541 |
| PAO1 3xFLAG-*pilJ* Δ*pilG* Δ*pilK* pUC18_PlacP1-YFP/POXB20-mKate2 | this study | RP582 |
| PAO1 3xFLAG-*pilJ* Δ*pilG* Δ*chpB* | this study | RP543 |
| PAO1 3xFLAG-*pilJ* Δ*pilG* Δ*chpB* pUC18_PlacP1-YFP/POXB20-mKate2 | this study | RP583 |
| PAO1 3xFLAG-*pilJ* Δ*pilH* | this study | RP514 |
| PAO1 3xFLAG-*pilJ* Δ*pilH* pUC18_PlacP1-YFP/POXB20-mKate2 | this study | RP529 |
| PAO1 3xFLAG-*pilJ* Δ*pilH* Δ*cyaB* | this study | RP569 |
| PAO1 3xFLAG-*pilJ* Δ*pilH* Δ*cyaB* pUC18_PlacP1-YFP/POXB20-mKate2 | this study | RP593 |
| PAO1 3xFLAG-*pilJ* $pilH_{D52A}$ | this study | RP730 |

| | | |
|---|---|---|
| PAO1 3xFLAG-*pilJ* *pilH*$_{D52A}$ pUC18_PlacP1-YFP/POXB20-mKate2 | this study | RP784 |
| PAO1 3xFLAG-*pilJ* *pilH*$_{D52A}$ Δ*cyaB* | this study | RP753 |
| PAO1 3xFLAG-*pilJ* *pilH*$_{D52A}$ Δ*cyaB* pUC18_PlacP1-YFP/POXB20-mKate2 | this study | RP793 |
| PAO1 3xFLAG-*pilJ* *pilH*$_{D52E}$ | this study | RP738 |
| PAO1 3xFLAG-*pilJ* *pilH*$_{D52E}$ pUC18_PlacP1-YFP/POXB20-mKate2 | this study | RP788 |
| PAO1 3xFLAG-*pilJ* *pilH*$_{D52E}$ Δ*cyaB* | this study | RP757 |
| PAO1 3xFLAG-*pilJ* *pilH*$_{D52E}$ Δ*cyaB* pUC18_PlacP1-YFP/POXB20-mKate2 | this study | RP795 |
| PAO1 3xFLAG-*pilJ* Δ*pilH* Δ*pilK* | this study | RP563 |
| PAO1 3xFLAG-*pilJ* Δ*pilH* Δ*pilK* pUC18_PlacP1-YFP/POXB20-mKate2 | this study | RP590 |
| PAO1 3xFLAG-*pilJ* Δ*pilH* Δ*chpB* | this study | RP545 |
| PAO1 3xFLAG-*pilJ* Δ*pilH* Δ*chpB* pUC18_PlacP1-YFP/POXB20-mKate2 | this study | RP584 |
| PAO1 3xFLAG-*pilJ* Δ*pilK* *pCuAIgent-empty* | this study | RP872 |
| PAO1 3xFLAG-*pilJ* Δ*pilK* *pCuAIgent-mNG-pilK* | this study | RP873 |
| PAO1 3xFLAG-*pilJ* Δ*pilK* Δ*chpB* *pCuAIgent-empty* | this study | RP874 |
| PAO1 3xFLAG-*pilJ* Δ*pilK* Δ*chpB* *pCuAIgent-mNG-pilK* | this study | RP875 |
| PAO1 Δ*pilG* pUCp19Δlac_PilG-HA (PA0408 C-terminus fused with HA tag) | Inclan *et al*, 2016[48] | JB1168 |
| PAO1 Δ*pilH* pUCp19Δlac_PilH-HA (PA0409 C-terminus fused with HA tag) | Inclan *et al*, 2016[48] | JB1171 |

**Key Resources Table 2: Plasmids used in this study.**

| **Name and relevant information** | **Source / Reference** | **Identifier** |
|---|---|---|
| pEX100TAP (Suicide vector based on pUC19, $Amp^R$, ColE1 ori (*E. coli*), *oriT*, *sacB*, *lacZα*) | Schweizer & Hoang, 1995[74] | Na |
| pEX18AP (Suicide vector based on pUC18, $Amp^R$, ColE1 ori (*E. coli*), *oriT*, *sacB*, *lacZα*) | Hoang *et al*, 1998[75] | Na |

| | | |
|---|---|---|
| pEX18GM (Suicide vector based on pUC18, $Gm^R$, ColE1 ori (*E. coli*), *oriT*, *sacB*, *lacZα*) | Hoang *et al*, 1998[75] | Na |
| pUC18_PlacP1-YFP/POXB20-mKate2 (fluorescent reporter for cAMP level: YFP controlled by the synthetic *LacP1* promoter and mKate2 controlled by POXB20 promoter (Oxford Genetics Ltd. (UK), Sigma) as reference. | Kühn *et al*, 2023[34] | YI996 |
| pEX100TAP-Δ*fliC* (PA1092) | Bertrand *et al*, 2010[30] | pJB215 |
| pEX100TAP-Δ*cpdA* (PA4969) | Inclan *et al*, 2011[36] | pJTW033 |
| pEX18GM-Δ*cpdA* (PA4969) | Kühn *et al*, 2021[33] | pMK019 |
| pEX100TAP-Δ*cyaB* (PA3217) | Inclan *et al*, 2011[36] | pJTW031 |
| pEX18GM-Δ*cyaB* (PA3217) | Kühn *et al*, 2021[33] | pMK018 |
| pEX100TAP-Δ*pilG* (PA0408) | Bertrand *et al*, 2010[30] | PJB118 |
| pEX100TAP-Δ*pilH* (PA0409) | Bertrand *et al*, 2010[30] | PJB119 |
| pEX18AP-Δ*pilGH* (PA0408 and PA0409 including intergenic region) | Kühn *et al*, 2023[34] | pXP322 |
| pEX18GM-$chpA_{LOF}$ (insertion of the histidine kinase domain of PA0413 with substituted residues D2086A, D2087A, G2088A) | Kühn *et al*, 2023[34] | pMK057 |
| pEX18AP-$pilH_{LOF}$ (PA0409 with substituted residue D52A) | Kühn *et al*, 2023[34] | pMK012 |
| pEX18AP-$pilH_{GOF}$ (PA0409 with substituted residue D52E) | Kühn *et al*, 2023[34] | pMK013 |
| pEX18AP-$pilG_{D58A}$ (PA0408 with substituted residue D58A) | Kühn *et al*, 2023[34] | pMK010 |
| pEX18AP-$pilG_{D58E}$ (PA0408 with substituted residue D58E) | Kühn *et al*, 2023[34] | pMK011 |
| tpEX18GM-*mNG-pilG* (PA0408 N-terminus fused with mNeonGreen, GGGGG linker) | Kühn *et al*, 2021[33] | YI883 |
| pEx100TAP-*mNG-pilH* (PA0409 N-terminus fused with mNeonGreen, GGGGG linker) | Kühn *et al*, 2021[33] | pXP125 |
| pEX18GM-*ΔpilG* for $pilH_{LOF}$ | this study | pMK014 |
| pEX18GM-*ΔpilG* for $pilH_{GOF}$ | this study | pMK015 |
| pEX18GM-*ΔpilK* (PA0412) | this study | RP123 |
| pEX18GM-Δ*chpB* (PA0414) | this study | RP124 |
| pEX18GM-$pilK_{LOF}$ (PA0412 with substituted residue R92E) | this study | RP376 |

| | | |
|---|---|---|
| pEX18GM-*chpB*$_{LOF}$ (PA0414 with substituted residue H184Y) | this study | RP365 |
| pEX18GM-*chpB-mNG* (PA0414 C-terminus fused with mNeonGreen, GGGGG linker) | this study | RP337 |
| pEX18GM-*chpB-mNG* (PA0414 C-terminus fused with mNeonGreen, GGGGS linker) | this study | pMK067 |
| pEX18GM-*mNG-pilK* (PA0412 N-terminus fused with mNeonGreen, GGGGG linker) | this study | RP333 |
| pEX18GM-*pilK-mNG* (PA0412 C-terminus fused with mNeonGreen, GGGGG linker) | this study | RP334 |
| pCuAIgent-empty (cumic acid-inducible plasmid, based on pFGL815 (Addgene 52322), empty insert can be removed by restriction digestion with XhoI and XbaI, $Gm^R$) | this study | pMK066 |
| pCuAIgent-*mNG-pilK* (PA0412 N-terminus fused with mNeonGreen, GGGGS linker, cumic acid inducible, $Gm^R$) | this study | pMK065 |
| pCuAI-*mNG-pilK* (PA0412 N-terminus fused with mNeonGreen, GGGGS linker, cumic acid inducible, $Kan^R$, template for pMK65) | this study | pMK035 |
| pEX18GM-3xFLAG-*pilJ* (PA0411 N-terminus fused with 3xFLAG tag, GGGGG linker) | this study | RP487 |
| pEX18GM-1xFLAG-*pilJ* (PA0411 N-terminus fused with 1xFLAG tag, GGGGG linker) | this study | YI928 |
| pUCpΔlac-*pilH*-HA (PA0409 C-terminus fused with HA tag) | (Inclan *et al*, 2016)[48] | pJB134 |
| pUCpΔlac-*pilG*-HA (PA0408 C-terminus fused with HA tag) | (Inclan *et al*, 2016)[48] | pJB132 |

#### Key Resources Table 3: Oligonucleotides used in this study.

| Identifier | Sequence | Purpose |
|---|---|---|
| oMK119 | CAT GCC TGC AGG TCG ACT GCA CGC TCG GCC TGT TCC | Generation of pMK14 and pMK15 |
| oMK120 | GGA AAC GGC CTG TTC CAT GTT CGC CCT ATA TCG AC | Generation of pMK14 and pMK15 |
| oMK121 | ATG GAA CAG GCC GTT TCC TGA TAT CCG GCC | Generation of pMK14 and pMK15 |

| | | |
|---|---|---|
| oMK122 | GCT ATG ACC ATG ATT ACG CCG CTC CAG CTC TGC ACC | Generation of pMK14 and pMK15 |
| oMK168 | CAT GCC TGC AGG TCG ACT GCA TAC CGA ACA TTA CCC GCG | Generation of pMK067 |
| oMK169 | TTC TTC ACC TTT AGA GAC CGA TCC ACC TCC ACC TGT TTC GAC TCC TGT CGG CG | Generation of pMK067 |
| oMK170 | GAT GAA TTG TAT AAA TAA ACA GGA GTC GAA ACA TGA ACC AG | Generation of pMK067 |
| oMK171 | GCT ATG ACC ATG ATT ACG ATA GAG GCT GGC GAA GCG | Generation of pMK067 |
| oMK014 | GTC TCT AAA GGT GAA GAA GAT AAT ATG G | Generation of pMK067 |
| oXP574 | TTA TTT ATA CAA TTC ATC CAT ACC CAT TAC | Generation of pMK067 |
| oMK113 | AAC CGC ACC AGG ACG GCC | Generation of pMK035 and pMK066 |
| oMK114 | GGT ATA TCT CCT TCT TAA AGT TAA ACA AAA TTA TTT CTA GTA ACG | Generation of pMK035 and pMK066 |
| oMK115 | CTT TAA GAA GGA GAT ATA CCA TGG TCT CTA AAG GTG AAG AAG ATA ATA TGG | Generation of pMK035 |
| oXP539 | TTT ATA CAA TTC ATC CAT ACC CAT TAC | Generation of pMK035 |
| oMK099 | ATG GAT GAA TTG TAT AAA GGT GGA GGT GGA TCG CAG GCG AAC GGC GTC TG | Generation of pMK035 |
| oMK116 | CTG GCC GTC CTG GTG CGG TTT CAT GTG CCT GAG TAC CCC TTA CG | Generation of pMK035 |
| oMK160 | TTA TTA TTT CCT TCC TCT TTT CTA CAG TAT TTA AAG ATA CC | Generation of pMK065 and pMK066 |
| oMK161 | TAC CTA GAA TGC ATG ACC AAA ATC CC | Generation of pMK065 and pMK066 |
| oMK162 | AAA GAG GAA GGA AAT AAT AAA TGT TAC GCA GCA GCA ACG ATG | Generation of pMK065 and pMK066 |
| oMK163 | TTG GTC ATG CAT TCT AGG TAT TAG GTG GCG GTA CTT GGG TC | Generation of pMK065 and pMK066 |
| oMK166 | CTT TAA GAA GGA GAT ATA CCT CGA GAC AGA GTA CAT CCT GCC CGC GTT TCT GGT CGG TTG | Generation of pMK066 (used as insert after annealing with oMK167) |

| | | |
|---|---|---|
| oMK167 | CTG GCC GTC CTG GTG CGG TTC TAG AGC GGC AGG CGT GTA ACA ACC GAC CAG AAA CGC GGG | Generation of pMK066 (used as insert after annealing with oMK166) |
| oRP94 | CGG GGA TCC TCT AGA CTG ACC GTG GCC GCG ACC | Generation of RP123 |
| oRP95 | CAC TCC ATT CCC CTC GTC CAA GGT GTC G | Generation of RP123 |
| oRP96 | ACG AGG GGA ATG GAG TGG CTA TGG GTG ACC | Generation of RP123 |
| oRP97 | GCC AGT GCC AAG CTT CCG CAC CAG TTC CTC GCA | Generation of RP123 |
| oRP119 | CGG GGA TCC TCT AGA GAG GTC AAG CAG CTC GGC GG | Generation of RP124 |
| oRP120 | TGC CTC GGC CGC AAC CCG TGG CGT GA | Generation of RP124 |
| oRP121 | GGT TGC GGC CGA GGC ATT GGT CAA GC | Generation of RP124 |
| oRP122 | GCC AGT GCC AAG CTT CAG GCG GCC GAG GAA CTC | Generation of RP124 |
| oRP165 | CGG GGA TCC TCT AGA CTG ACC GTG GCC GCG ACC | Generation of RP376 |
| oRP166 | GCC AGT GCC AAG CTT CCG CAC CAG TTC CTC GCA C | Generation of RP376 |
| oRP239 | CCA GGA AAC CCG TTT CTT CGA GCA TCC CCC GTC TTT CG | Generation of RP376 (using product of oRP165 and oRP166 as template) |
| oRP240 | CGA AAG ACG GGG GAT GCT CGA AGA AAC GGG TTT CCT GG | Generation of RP376 (using product of oRP165 and oRP166 as template) |
| oRP167 | CGG GGA TCC TCT AGA GAG GTC AAG CAG CTC GGC | Generation of RP365 |
| oRP168 | GCC AGT GCC AAG CTT CAG GCG GCC GAG GAA CTC | Generation of RP365 |
| oRP231 | CTT CCT CTA TGC CCA GTA CAT CGA TGC CAG CTT | Generation of RP65 (using product of oRP167 and oRP168 as template) |

| | | |
|---|---|---|
| oRP232 | AAG CTG GCA TCG ATG TAC TGG GCA TAG AGG AAG | Generation of RP65 (using product of oRP167 and oRP168 as template) |
| oRP175 | CGG GGA TCC TCT AGA GTT GCG GTG ATC GCC GAC | Generation of RP337 |
| oRP176 | CCG CCT CCT TCG ACT CCT GTC GGC GC | Generation of RP337 |
| oRP177 | AGT CGA AGG AGG CGG AGG CGG AAT G | Generation of RP337 |
| oRP178 | GCC TGG TTC ATG TTT TAT ACA ATT CAT CCA TAC CCA TTA CAT CAG | Generation of RP337 |
| oRP179 | TGT ATA AAA CAT GAA CCA GGC CGT GAT CG | Generation of RP337 |
| oRP180 | GCC AGT GCC AAG CTT AGC CGG GCG CCC AGG ATC | Generation of RP337 |
| oRP181 | CGG GGA TCC TCT AGA GTG GCC GCG ACC GTG ACC | Generation of RP333 |
| oRP182 | GAG ACC ATG CCG TGC CCC TCG TCC AAG | Generation of RP333 |
| oRP183 | GCA CGG CAT GGT CTC TAA AGG TGA AGA AGA TAA TAT GG | Generation of RP333 |
| oRP184 | CCT GCA TTC CGC CTC CGC CTC CTT TAT AC | Generation of RP333 |
| oRP185 | GAG GCG GAA TGC AGG CGA ACG GCG TC | Generation of RP333 |
| oRP186 | GCC AGT GCC AAG CTT TCG GGT CCT GCG GGT TCT C | Generation of RP333 |
| oRP187 | CGG GGA TCC TCT AGA AAG ATG GCG AGC GAG ATG | Generation of RP334 |
| oRP188 | CCG CCT CCT GTG CCT GAG TAC CCC TTA C | Generation of RP334 |
| oRP189 | AGG CAC AGG AGG CGG AGG CGG AAT G | Generation of RP334 |
| oRP190 | CCA CTC CAT TCA TTT ATA CAA TTC ATC CAT ACC CAT TAC ATC AG | Generation of RP334 |
| oRP191 | TGT ATA AAT GAA TGG AGT GGC TAT GG | Generation of RP334 |
| oRP192 | GCC AGT GCC AAG CTT CAC CAG TTC CTC GCA CAAG | Generation of RP334 |

| | | |
|---|---|---|
| oRP282 | CGA CTA CAA AGA CGA TGA CAA AAT CGA TGA CTA CAA AGA CGA TGA CAA AGG CGG CGG CGG CGG CAT GAA GAA AAT CAA CG | Generation of RP487<br>(using YI928 as template) |
| oRP283 | TTT TGT CAT CGT CTT TGT AGT CGG CGG CTT TGT CAT CGT CTT TGT AGT CCA TAT TTG GCC CCC GCC CGG ACC GAC CAC GA | Generation of RP487<br>(using YI928 as template) |
